# Mapping spatially-resolved transcriptomes in systemic sclerosis

**DOI:** 10.1101/2025.01.14.632962

**Authors:** Zhijian Li, Aleix Rius Rigau, Wenjie Xie, Linlin Huang, Xiaohang Shao, Yi-Nan Li, Alexandru-Emil Matei, Wenjing Ye, Hejian Zou, Luca Pinello, Jörg H.W. Distler, Rui He, Minrui Liang

## Abstract

Systemic sclerosis (SSc) is a prototypical fibrotic disease with high mortality and limited treatment options. Despite advances in single-cell RNA sequencing (scRNA-seq), the comprehensive understanding of cellular heterogeneity and cell-cell interaction within the fibrogenesis microenvironment remains limited. We generated spatially resolved transcriptome maps from healthy and SSc skin and built a scRNA-seq atlas to map the single-cell data to spatial space. This enabled us to identify a fibrotic niche, enriched with fibroblasts and macrophages, which is significantly expanded in SSc and correlated with clinical outcome. We revealed disease-specific cell states of fibroblasts and macrophages, and evaluated their spatial dependency on other cell types. We identified selective expression of ACKR3 in fibroblast progenitors that diminishes with SSc progression, which may serve to regulate CXCL12/CXCR4-mediated macrophage recruitment and fibrotic remodeling. Together, we provided an in-depth description at cellular and spatial levels of fine-tuned regulatory events occurring in SSc, offering spatiotemporal insights.

**One Sentence Summary:** Integrated spatial omics provide insight into the cellular and transcriptional landscape in spatially distinct microenvironments, which may drive fibrosis progression in SSc.

## INTRODUCTION

Systemic Sclerosis (SSc), an orphan autoimmune disease, is characterized by the progressive fibrosis affecting skin and multiple organs, holding the highest mortality among all rheumatic diseases(*1*). The development of effective therapeutic strategies for SSc is impeded by a lack of comprehensive understanding of its pathogenesis and the limited translatability of current preclinical models. Importantly, studying SSc as a prototypical systemic fibrotic disease could yield new insights into the pathogenesis of fibrotic diseases, and aid in the discovery of novel therapeutic approaches.

SSc is characterized by a complex interplay of features that include vasculopathy, inflammation, and fibrosis. Within this triad, fibroblasts have been recognized as pivotal effector cells in the production of extracellular matrix (ECM), largely contributing to developing fibrotic tissue remodeling in SSc. Fibroblasts sense and receive signaling from the neighboring cells, which are conveyed and converge to the shared and distinct profibrotic pathways(*2*). The coordination of fibroblasts with macrophages can shape their cellular response, therefore orchestrating the initiation or resolution of fibrosis across tissues(*3–7*). Beyond the conventional macrophage polarization paradigm, new populations of macrophages have been linked with tissue repair and fibrotic remodeling(*3–5*, *7*, *8*). For instance, phenotypically distinct macrophages (Lyve1^lo^MHCII^hi^ and Lyve1^hi^MHCII^lo^) manifested specific niche-dependent functional roles in pulmonary and cardiac fibrosis(*8*). However, it remains largely unexplored whether and how the spatial neighbourhood in the tissue architecture affects the crosstalk between fibroblast and macrophage subsets, potentially driving fibrotic remodeling in SSc.

Recent studies using single-cell RNA sequencing (scRNA-seq) have provided an unprecedented opportunity to study the molecular mechanisms of SSc at single-cell resolution(*9–13*). Pioneering studies have characterized fibroblast types and states in SSc(*9*, *11*, *13*). While these studies have provided valuable insights, they are significantly constrained by the lack of spatial resolution due to the reliance on disaggregated tissue analysis. We recently demonstrated cellular niches centered around endothelial cells or fibroblasts in SSc using a panel of up to 40 markers through multiplex spatial proteomics(*14*, *15*). Thus, a spatially-resolved characterization of genome-wide transcriptomics of SSc is needed.

In this work, we profiled human skin from SSc patients and healthy controls using spatial transcriptomics (ST) technique based on the 10x Visium formalin-fixed paraffin-embedded (FFPE) platform and spatial enhanced resolution omics sequencing (Stereo-seq)(*16*). To increase the cellular resolution of our ST data, we integrated two publicly available scRNA-seq datasets, creating a human skin transcriptome atlas and mapping cell type information onto spatial regions. We validated our results using additional spatial profiling techniques with higher resolution, such as imaging mass cytometry (IMC), as well as time-course experimental skin fibrosis. These approaches enabled us to reveal a detailed characterization of pathological cellular communication between fibroblast and macrophage subsets. This has provided insights into therapeutic target discovery by disrupting disease-driving cellular niches and circuits in SSc.

## RESULTS

### Profiling of SSc and healthy skin with spatially-resolved transcriptomics

We collected a total of 14 human skin biopsies from 10 SSc patients enriched for early-stage, diffuse cutaneous involvement, with clinically active disease and 4 sex- and age-matched normal controls (**Fig. 1A and table S1**). For each biopsy, we generated spatially resolved transcriptomics (ST) using the 10x Visium platform. After removing low-quality spots across all samples, we retained 13,204 high quality spots with a median detection of 2,265 genes per spot (**figs. S1, A to D, and table S2**). To enhance spatial resolution, we also performed high-resolution ST profiles on four samples, one from a healthy donor and three from SSc patients, using Stereo-seq. After controlling data quality, we retained 40,566 spots across these samples. We additionally analyzed two scRNA-seq datasets generated from skin tissues of SSc patients (n = 104) and healthy donors (n = 64)(*9*, *10*). The cells were re-annotated for each dataset based on clustering or the original labels (**figs. S2, A to D; see Materials and Methods**). We integrated the scRNA-seq datasets and observed that the same cell types were well-aligned in the UMAP space, which was further confirmed by the cell-type-specific marker genes identified based on the integrated profiles (**Figs. 1, B to C, figs. S2, E to G**). Together, these data provide a comprehensive multi-modal map of human skin tissues in health and SSc, capturing both spatial and single-cell transcriptomic landscapes.

**Fig. 1.**
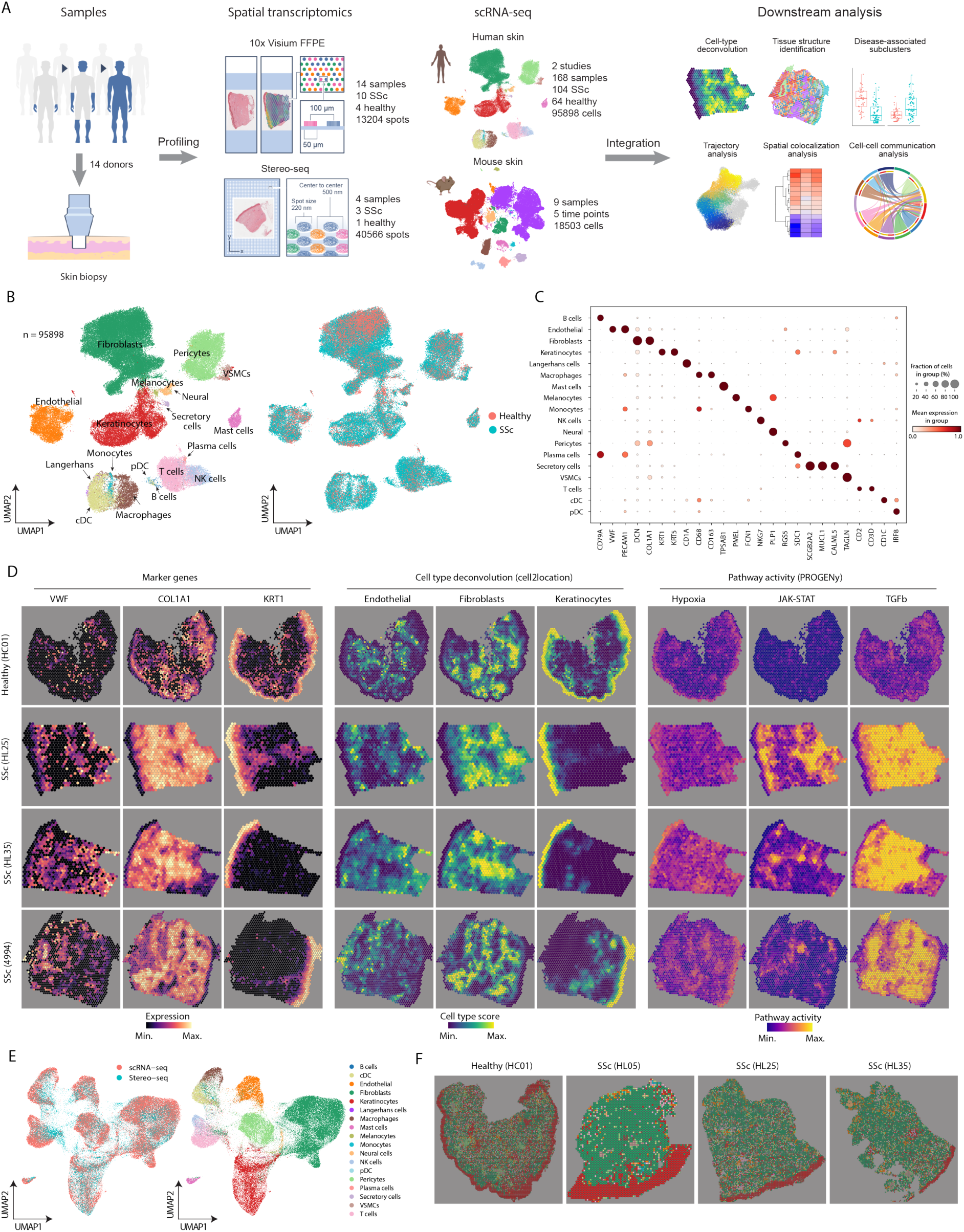
Spatial profiling of human skin. (**A**), Overview of experimental design and analysis workflow. (**B**), UMAP showing integrated scRNA-seq datasets as colored by cell types (left) and conditions (right). SSc skin n=104, Healthy skin n=64. (**C**), Dotplot showing gene expression for marker genes. The colors refer to the mean expression of the genes and the dot sizes represent the proportion of cells expressing the marker genes. (**D**), Visualization maps based on 10× Visium. SSc skin n=10, Healthy skin n=4. Left: *vWF*, *COL1A1*, and *KRT1* gene expressions in spatial transcriptomics data. Middle: representative cell types estimated by cell2location. Right: visualization of pathway activity in spatial space. (**E**), Integration of scRNA-seq and Stereo-seq data colored by protocols (left) and cell types (right). (**F**), Visualization of cell types in spatial space based on Stereo-seq data. SSc skin n=3, Healthy skin n=1. The experiments were performed on adjacent tissue sections to those used for 10× Visium.

We next deconvoluted the cell-type compositions of each spot in 10x Visium data using the scRNA-seq as reference. The estimated cell type abundances from spatial transcriptomics were generally co-localized with their respective marker genes (**Fig. 1D**). Moreover, we observed that different cell types were mapped to the expected locations. For example, keratinocytes were found localized within the epidermis and surrounding the hair follicle. Endothelial cells were closely associated with vasculature in the dermis. Fibroblasts were distributed throughout the papillary layer and reticular layer of the dermis. We inferred the pathway activity for each spot and observed the enriched expression of genes associated with some well-known pathways, such as Janus kinase signal transducer and activator of transcription protein (JAK-STAT) and transforming growth factor (TGF)-β, reflecting the link between cellular location and the corresponding function (**Fig. 1D**). We transferred the cell type annotation from scRNA-seq to Stereo-seq through data integration (**Fig. 1E**). Spatial visualization of the predicted labels in tissues showed a similar cellular distribution as 10x Visium data (**Fig. 1F**). Collectively, our integrative analyses on spatial transcriptomics and scRNA-seq data enable us to map cell abundance and signaling pathways across zonations in skin samples.

### Characterization of tissue organization in SSc and healthy skin

To characterize the cellular colocalization events in human skin, we integrated the 10x Visium samples (n = 14) based on the inferred cell-type proportions within each spot (**fig. S3A**). Specifically, we conducted spot deconvolution to determine the proportions of different cell types within each spot, and subsequently clustered the spots based on these identified proportions (**Fig. 2A**). Clustering analysis identified 12 compositionally distinct populations, defined as cell-type niches that represent potential structural building blocks shared across different samples. Visualization of the cell-type abundances and marker genes of these niches revealed that they were strongly associated with the corresponding cell types (**Fig. 2B, figs. S3, B to C**). For example, niches 1 and 3 (defined as vasculature niches) were enriched with endothelial cells and pericytes; niches 0 and 10 (defined as fibrotic niches) were enriched with fibroblasts; niches 6, 7 and 8 (defined as epidermis niches) were largely associated with keratinocyte as the representative cell types and along with the infiltration of Langerhans cells; niche 11 was related to vascular smooth muscle cells (VSMC) and therefore defined as a VSMC niche; niche 2, 4, 5, and 9 (defined as immune niches) were related to immune cells (**Fig. 2C**).

**Fig. 2.**
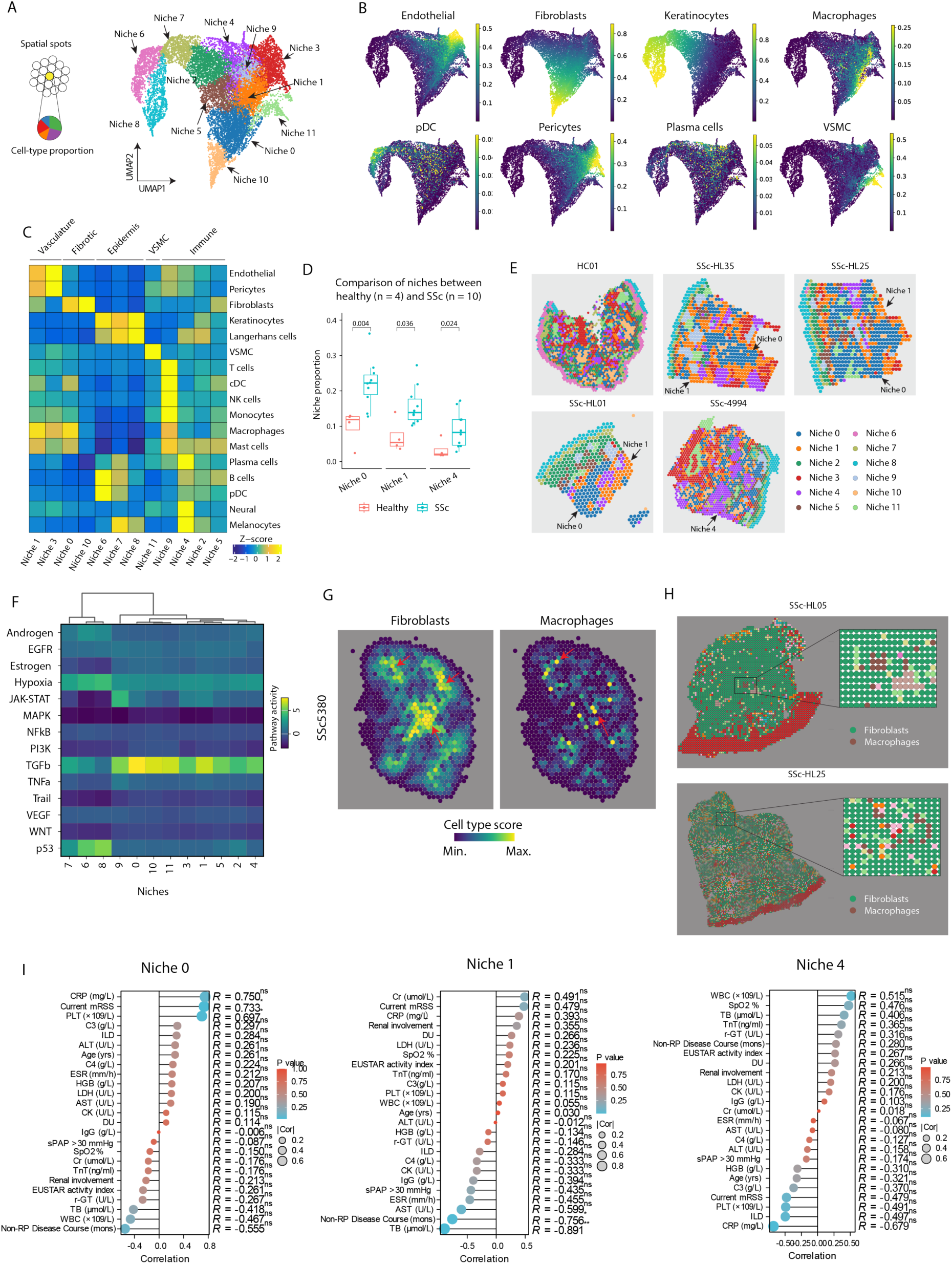
Characterization of cell-type niches in SSc and healthy skin. (**A**), UMAP showing the integrated spatial data as colored by cell-type niches. (**B**), Visualization of cell type proportion in UMAP space. (**C**), Heatmap showing the enrichment of cell types in niches. Each column represents a niche, and each row represents a cell type. Colors refer to the z-score transformed proportions of each cell type across different niches. (**D**), Boxplot comparing the proportion of each niche between SSc and healthy skin samples. Each dot represents a sample. *P*-values are calculated using the Wilcoxon rank-sum test (unpaired; two sides). (**E**), Visualization of cell-type niches in spatial space for the healthy and SSc samples using 10x Visium data. (**F**), Heatmap showing pathway activity for each niche. (**G**), Visualization of fibroblasts and macrophages in spatial space using 10x Visium data. (**H**), Visualization of cell types in spatial space using Stereo-seq data. (**I**), Correlations between the proportion of the indicated niches with clinic statistics across all SSc subjects. Statistical significance was determined by the Spearman test for a two-variable correlation. The correlation coefficient r is indicated in figures (* *p* < 0.05).

Furthermore, we observed a significant increase in the proportions of three niches (0, 1, and 4) in the SSc skin compared to healthy skin (**Figs. 2, D to E, fig. S3D**). Notably, niche 0 was infiltrated with a high prevalence of fibroblasts, macrophages, conventional dendritic cells (cDCs), and mast cells, indicating a potential immune microenvironment within a fibrogenic niche. Niche 1 was characterized by an enrichment of endothelial cells, pericytes, macrophages, cDCs, and mast cells, suggesting a perivascular niche associated with immune cell infiltration in SSc skin. Additionally, niche 4 was distinguished by a significant presence of plasma cells, melanocytes, and neural cells (**Fig. 2C**). Moreover, SSc skin exhibited a more reorganized structure, highlighting a tendency towards spatially educated organization, with a dominance in niche distribution (**Fig. 2E, fig. S3D**).

To investigate the featured signaling pathway that characterizes each specific niche, we computed pathway activity. Fibroblast-dominant niche 0 showed the highest activity of the TGF-β signaling pathway. Niche 9, composed of various immune cells, displayed the strongest activity of JAK-STAT signaling (**Fig. 2F**). These findings supported the essential roles of TGF-β and JAK-STAT signaling in the pathogenesis of SSc as expected(*2*), suggesting a convergence of multiple signaling activities within the structural organization in SSc fibrosis. KEGG pathway enrichment analysis based on marker genes also revealed high activity in the TGF-β signaling in niche 0 (**fig. S3E**).

Indeed, we observed colocalization of fibroblasts and macrophages in our 10x Visium and Stereo-seq data (**Figs. 2, G to H**). To further validate these results, we performed imaging mass cytometry (IMC) and identified the cellular neighborhood (CN) community as described previously(*14*). We observed similar niches comprising stromal cells and macrophages (CN8), vascular ECs, and immune cells, including macrophages in CN5 (**figs. S4, A to C**). To explore the clinical relevance of these niches, we correlated their proportions with clinical data across the SSc samples. Interestingly, we observed a positive Spearman correlation between the proportion of niche 0 and the current total modified Rodnan’s skin score (mRSS) (*r* = 0.733, *p* = 0.023), C-reactive protein (CRP) levels (*r* = 0.750, *p* = 0.066) and platelet counts (*r* = 0.697, *p* = 0.031), which are interpreted as clinical indicators for disease severity or activity of SSc (**Fig. 2I**). Additionally, we observed a significant inverse correlation between the proportion of niche 1 and the non-Raynaud’s phenomenon (RP) disease duration (*r* = -0.756, *p* = 0.014). In summary, our spatial multi-omics revealed a distinct structural organization associated with disease outcome of SSc, suggesting a potential coordinated regulation of immune-stromal interactions that may drive the disease progression.

### Characterization of fibroblast sub-clusters in healthy and SSc skin

To investigate the molecular mechanisms driving fibrogenesis in human SSc, we next clustered scRNA-seq data for fibroblasts and identified ten sub-clusters across the datasets and conditions (**Fig. 3A; fig. S5A**). Consistent with previous studies(*9*, *10*), we found the corresponding fibroblast populations, including *SCARA5^+^*/*PCOLCE2^+^/LGR5^+^*fibroblasts (Fib1), *MYOC^+^/C3^+^/APOD^+^* fibroblasts (Fib3), *CCL19^+^/C7^+^/APOE^+^* fibroblasts (Fib4), *SFPR2^+^/WIF1/APCDD1^+^* fibroblasts (Fib6), *COCH^+^/SPARCL1^+^*fibroblasts (Fib7), *COMP*^+^/*SFPR2^+^*/*PRSS23^+^*fibroblasts (Fib9), and *IGFBP2^+^/ANGPTL7^+^* fibroblasts (Fib10) (**Fig. 3B**). Moreover, Fib5 was characterized by the expression of *FOS*, *JUNB*, *JUN* and *FOSB* which were components of activator protein 1 (AP1) known as the core transcription factors in SSc fibrosis(*17*, *18*). Fib1 demonstrated the upregulation of *SCARA5*, which has been recently identified as a marker for progenitor fibroblast in human myocardial and kidney fibrosis(*19–21*). Fib2 and Fib9 were characterized by the expression of Periostin (*POSTN*), which encodes an extracellular matrix protein that can amplify TGF-β signaling(*22*, *23*). In addition, Fib2 also exhibited a high expression of myofibroblast hallmark *ACTA2* (**Fig. 3B**). Together, these suggested that Fib2 and Fib9 as terminally-differentiated ECM-producing myofibroblasts.

**Fig. 3.**
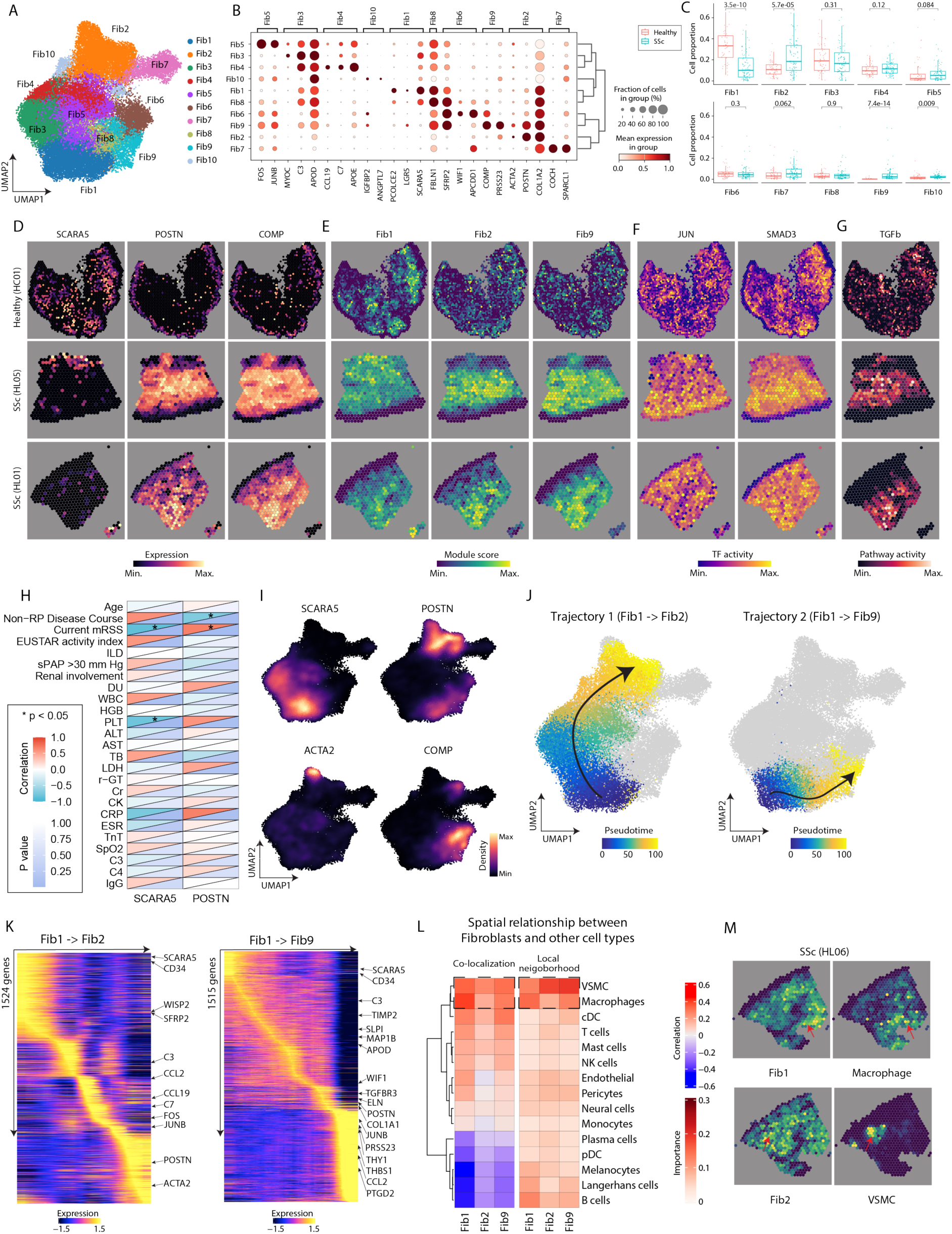
Characterization of fibroblast sub-populations. (**A**), UMAP showing sub-populations of fibroblasts based on scRNA-seq data. (**B**), Dot plot showing the marker genes of each subset. (**C**), Bar graph comparing the proportion of each fibroblast subset between healthy and SSc samples. Each dot represents a sample. *p*-values are calculated using the Wilcoxon rank-sum test (unpaired; two sides). (**D**), Visualization of *SCARA5, POSTN*, and *COMP* expression in spatial space. (**E**), Visualization of module score of Fib1, Fib2, and Fib9 in spatial space. (**F**), Visualization of TF activity of JUN and SMAD3 in spatial space. (**G**), Visualization of TGF-β pathway activity in spatial space. (**H**), Heatmap showing correlations between expressions of *SCARA5* and *POSTN* and different clinical outcomes of SSc. The colors in the upper left square represent Spearman’s r value, while those in the lower right square represent p values (* *p* < 0.05). (**I**), Expression of *SCARA5, POSTN, ACTA2*, and *COMP* overlaying with UMAP. (**J**), UMAP showing the differentiation trajectory from Fib1 to Fib2 (left), and Fib1 to Fib9 (right). Colors refer to pseudotime points. (**K**), Heatmap showing the gene expression dynamics during the trajectory from Fib1 to Fib2 (left), and Fib1 to Fib9 (right). (**L**), co-localization between major cell types and fibroblast sub-clusters (Fib1, Fib2 and Fib9) as measured within the same spot (left) and neighboring spots (right). (**M**), Visualization of the abundance of macrophages and VSMCs and the state scores of Fib1 and Fib2 in a representative SSc skin sample.

Next, we compared the proportion of each sub-cluster between healthy and SSc samples. We found that Fib1 showed a significant reduction in SSs skin (average fraction = 0.137) compared to healthy skin (average fraction = 0.319) (*p* = 3.5e-10). In contrast, Fib2 exhibited an increase in SSc skin (average fraction = 0.224) compared with that in healthy skin (average fraction = 0.114) (*p* = 5.7e-05) (**Fig. 3C**). We also estimated transcription factor (TF) activities for each subset and observed that Fib2 was characterized by high TF activity of the hedgehog transcription factor GLI2 (**fib. S5B**). This convergent driver can mediate Hedgehog pathways in activating fibroblast and promoting fibrosis in SSc(*18*, *24*). Moreover, Fib9 was marked by *SMAD3*, *SMAD4*, and *JUN*, all of which are TGF-β downstream pathways to promote fibrosis. To validate our results, we re-analyzed a bulk RNA-seq dataset from *PRESS* cohort(*25*) and observed the significant increase of *POSTN* and decrease of *SCARA5* in SSc skin compared with healthy skin (**fib. S5C**). An inverse correlation was also observed in the gene expressions between *SCARA5* and *POSTN* (**fig. S5C**). Overall, these findings further supported the potential involvement of *SCARA5*^+^ and *POSTN*^+^ fibroblasts in the pathogenesis of SSc.

We next extended our analysis to spatial transcriptomics data. Visualization and quantification of gene expression showed that *SCARA5* was significantly downregulated in SSc skin, whereas *POSTN* was upregulated in SSc skin (**fig. S5D**). Additionally, *SCARA5* has exhibited distinct spatial enrichment patterns that were mutually exclusive to regions enriched for *POSTN* and *COMP* across multiple samples (**Fig. 3D and fig. S5E**). To characterize the spatial distribution of fibroblast subsets, we calculated module scores of each subset, defined as the average expression of the top 100 marker genes, across spatial spots based on their marker genes. This analysis revealed that Fib2 and Fib9 were spatially colocalized (**Fig. 3E and fig. S5F**). To probe the regulatory networks underlying these subsets, we estimated the activity of TFs and signaling pathways, and observed increased activities of JUN, SMAD3, and the TGF-β pathway, which were closely associated with the spatial localization of Fib2 and Fib9. These findings align with their known pro-fibrotic roles, further confirming the spatial mapping of these sub-clusters (**Figs. 3, F and G, figs. S5, G and H**). To explore the clinical relevance of these findings, we correlated *SCARA5* and *POSTN* expression with clinical outcomes. Interestingly, *SCARA5* expression negatively correlated with the mRSS, whereas *POSTN* expression positively correlated with mRSS (**Fig. 3H**). Additionally, *SCARA5* showed an inverse correlation with platelet counts, while *POSTN* negatively correlated with non-RP disease duration. These results collectively suggested divergent effects of *SCARA5*^+^ and *POSTN^+^* fibroblasts in SSc fibrosis.

### Differentiation of *SCARA5^+^* to *POSTN^+^* fibroblasts in SSc skin

While pioneering studies have identified *SCARA5*^+^ fibroblasts as myofibroblast progenitors in fibrosis^19–21^, their differentiation trajectories have not been constructed yet. To firstly determine if this phenomenon is consistent across different tissues such as SSc skin fibrosis, we analyzed scRNA-seq data from mouse skin challenged by bleomycin (BLM) at various time points (days 3, 7, 14, and 28) and compared them to untreated skin (day 0, steady state)(*26*). This analysis identified 14 distinct cell types and nine fibroblast sub-clusters (**figs. S6, A and B**). Among these, we found a progressive decrease in *Scara5*^+^ fibroblasts following BLM treatment, while *Postn*^+^ fibroblasts substantially increased on days 14 and 28 (**figs. S6, C and D**). These temporal changes suggested a potential differentiation process from *Scara5*^+^ to *Postn*^+^ fibroblasts.

Based on these observations and the fact that Fib2 and Fib9 were marked by *POSTN*, we next inferred two pseudo-time trajectories from Fib1 to Fib2 (trajectory 1) and Fib1 to Fib9 (trajectory 2) (**Fig3. I and J**). To characterize transcriptional dynamics during the differentiation, we estimated the expression variation of each gene along the pseudo time and identified 1524 and 1515 genes associated with trajectory 1 and 2, respectively. For trajectory 1, we found many genes were down-regulated during the differentiation, including precursor marker *CD34*(*27*) and *SFRP2*(*9*) (**Fig. 3K, fig. S6E**). Moreover, we identified a set of genes, including *C3*, *C7*, *CCL2*, and *CCL19*, that were particularly enriched at the intermediate stage of the trajectory. This suggested a role for chemokine and cytokine signaling, as well as complement system activity, in the differentiation process occurring at this stage. Genes associated with the late stage of this trajectory include *FOS* and *JUNB,* which are involved in the fibrosis-modulatory pathway and have been observed to be activated during the myofibroblast transition process(*2*). For trajectory 2, we additionally observed that *TIMP2, TGFBR3* and *APOD* were associated with the intermediate stage (**Fig. 3K, fig. S6F**). These results revealed distinct gene regulatory networks that are involved in the divergent differentiation of fibroblasts.

Of note, Sarthak Sinha et al conducted comparative single-cell multi-omics in reindeer following full-thickness injury and found that the velvet skin regenerates, while the back skin heals by forming a scar(*7*). Similarly, in reindeer skin, *SCARA5* gradually decreased but *ACTA2* increased post wound in their back, but not in velvet (**figs. S7, A to E**). This pattern is reminiscent of fibrotic tissue remodeling in human, which indicates the fibroblast dynamic changes could be conserved across species and tissues.

### Spatial co-localization of fibroblast sub-clusters and other cell types

Apart from highly discordant fibroblast states, the distinct stromal-immune linkages also determine the fibrotic progression. Next, we investigated spatial dependencies between fibroblast subsets and other major cell types to better understand their spatial communications and contributions to fibrotic microenvironments. We conducted spatial colocalization analyses in two distinct proximity areas: 1) co-occurrence within one single spot, determined by Spearman correlation between the module score of each fibroblast sub-cluster and the proportion of each major cell type, and 2) extended neighbourhood within an annular region encompassing up to three surrounding spots for each target spot analyzed with the MlSTy pipeline(*28*). The results from both approaches were averaged across all samples to identify consistent spatial patterns. We focused specifically on Fib1, Fib2, and Fib9, as they have been demonstrated to exhibit significant alterations in SSc (**Fig. 3C**). Notably, we observed a consistent spatial dependency in both models across tissues, characterized by an increased co-occurrence between fibroblast progenitor *SCARA5*^+^ Fib1 and macrophages, whereas *POSTN*^+^/*ACTA2*^+^ myofibroblasts (Fib2 and Fib9) were found in proximity to VSMCs (**Figs. 3, L and M**). These findings point towards the potential critical role of macrophage in establishing the spatial microenvironment surrounding fibroblast from the early stage.

### Correlations of the cellular phenotypes with clinical outcomes of SSc

Next, we analyzed the potential correlations between the cellular phenotypes of fibroblast subpopulations and their respective niches with the clinical outcomes and treatment responsiveness in patients with SSc. The proportion of niche 0, which were predominantly composed of fibroblasts and macrophages, were significantly higher in the SSc patients who were non-responsive to a 52-week standard treatment of Mycophenolate mofetil (MMF), compared to those who were responsive (median proportion in responder is 18.28%, n=5; median proportion in non-responder is 25.00%, n=5, *p*=0.0079, determined by two-side Wilcoxon rank sum test) (**Fig. 4A**). All the SSc patients included in our study have been under the standard treatment of MMF for consecutive 52 weeks. The responsiveness was defined according to the criteria of the revised ACR-CRISS (American College of Rheumatology Composite Response Index in Systemic Sclerosis), which is the improvement in ≥ 3 of 5 core clinical measurements with Minimal Clinically Important Difference (MCID) of 25% for modified Rodnan skin score (mRSS), 5% for percent predicted forced vital capacity (FVC%), 20% for health assessment questionnaire-disability index (HAQ-DI), 20% for patient (PGA) and clinician (CGA) global assessments(*29*). Consistently, the proportions of niche 0, which is enriched with fibroblasts and characterized by high activity of TGF-β signaling, were correlated negatively with *SCARA5*, and positively with *POSTN*, *ACTA2*, more pronounced with the gene expression ratios of *POSTN/SCARA5*, and *ACTA2/SCARA5* (**Figs. 4, B to E**). Moreover, the largely overlapping of Niche 0 and *POSTN^+^/ACTA2^+^* myofibroblasts (Fib2 and Fib9), as quantified by module scores based on the average expression of the top 100 marker genes, which was observed by visualization using the ST data in SSc skin (**Fig. 4D**), which indicates niche 0 was featured by the molecular phenotype of myofibroblast rather than fibroblast progenitor. Furthermore, the ratios of *POSTN/SCARA5* and *ACTA2/SCARA5* manifested the ability to discriminate the SSc patients from healthy controls with the AUC of 0.875 and 0.800 (*p*=0.0339 and 0.0897, respectively, **Fig. 4F**). Those baseline ratios were observed to be significantly increased in the patients who experienced skin fibrosis progression (**Fig. 4G**), determined as over 25% increment in mRSS assessed at 52-week follow-up. Notably, the higher baseline ratios of *POSTN/SCARA5* and *ACTA2/SCARA5* were closely linked to the non-responsiveness for the standard treatment with MMF for 52 weeks (**Fig. 4H**). These data highlighted the potentials of aforementioned gene expression ratios in identification of the standard-treatment-refractory SSc patients, who may require more advanced treatment regimens rather than standard MMF. This provides insights into the future precision stratification strategy for SSc patients.

**Fig. 4.**
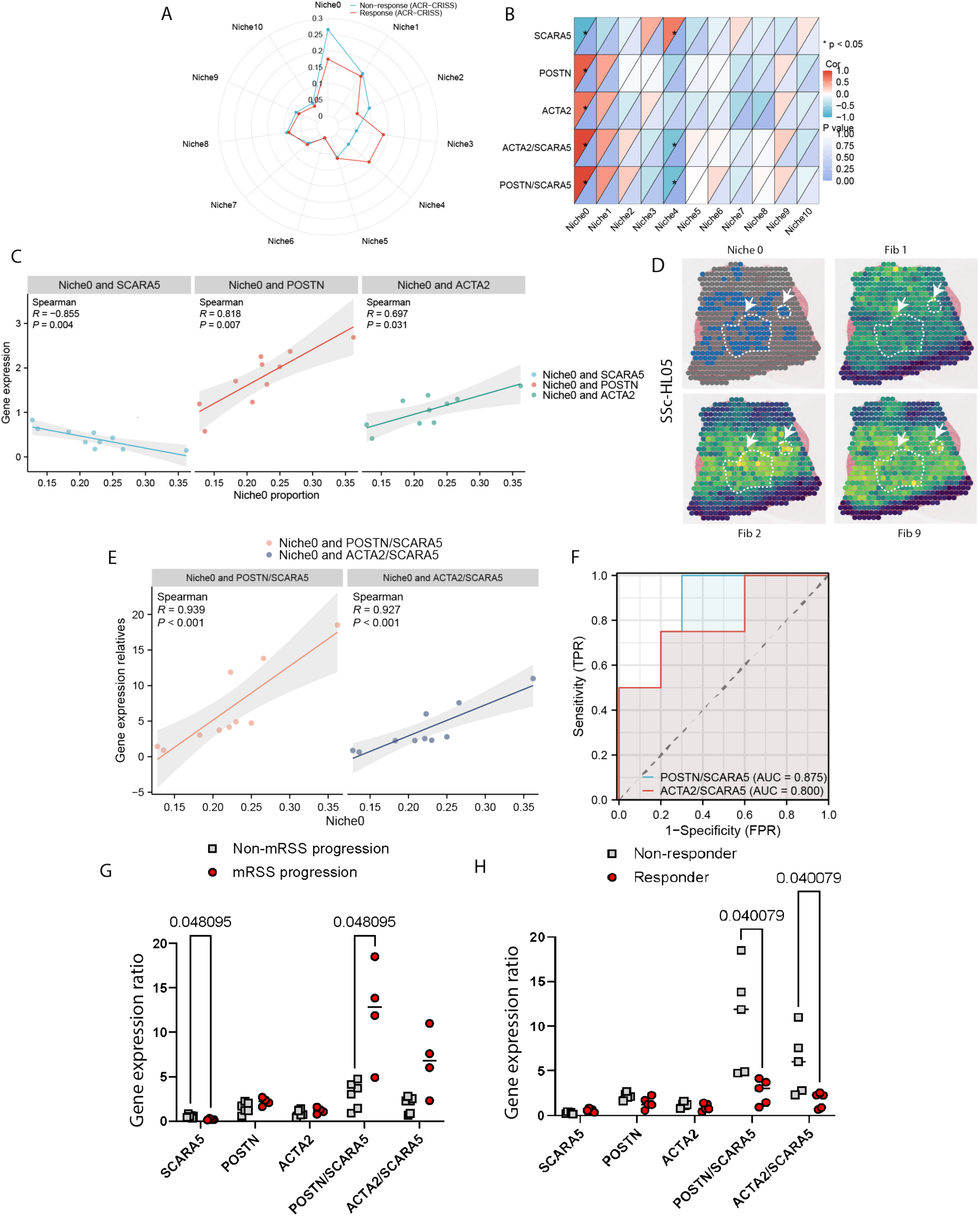
Correlations of the cellular phenotype with clinical outcomes. (**A**), A radar plot displaying the proportions of the identified niches in SSc patients, distinguishing between those who were responsive to the standard treatment of Mycophenolate Mofetil (MMF) and those who were not. (**B**), Heatmap showing correlations between the proportion of each niche and the pseudo-bulk gene expression, along with the ratios of *POSTN/SCARA5* and *ACTA2/SCARA5*. The colors in the upper left square represent Spearman’s r value, while those in the lower right square represent *p* values (* *p* < 0.05). (**C**), Correlations between the proportion of niche 0, which was predominantly composed of fibroblasts and macrophages, and the gene expressions of *SCARA5, POSTN, ACTA2*, along with their ratios. Correlation data were analyzed using non-parametric Spearman tests. Each data point represents one SSc sample, and the confidence interval (CI) is indicated by grey shadows. (**D**), Visualization of the abundance of niche 0 and the state scores of Fib1, Fib2 and Fib9 in a representative SSc skin sample. (**E**), Receiver operating characteristic (ROC) curve showing the discrimination of SSc patients from healthy controls based on the gene expression ratios of *POSTN/SCARA5* and *ACTA2/SCARA5*, respectively. The area under the curve (AUC) values are indicated in graphs. (**F**), The gene expressions of *SCARA5, POSTN, ACTA2*, along with their ratios, comparing the SSc patients who experienced skin fibrosis progression or not. (**G**), The gene expression of *SCARA5, POSTN, ACTA2*, along with their ratios, comparing the SSc patients who were responsive to the standard treatment of MMF or not, evaluated at a 52-week follow-up. Two-sided Wilcoxon rank sum tests were conducted to compare the differences between the two groups.

### Characterization of macrophage heterogeneity

We therefore asked how the immune microenvironment shapes the differentiation and phenotypes of fibroblasts, and vice versa. Beyond the conventional polarization program of macrophage, new subpopulations have been implicated in tissue fibrosis(*3–5*, *30*). They are not the direct source of ECM, but rather a major source of immune dysregulation in fibrotic conditions through release of proinflammatory and profibrotic factors(*31*). Nevertheless, the heterogeneity of macrophages and their crosstalk with mesenchymal cells in the diseased context of SSc remain poorly defined. We thus clustered the scRNA-seq data of macrophages and identified three transcriptionally distinct subsets (**Fig. 5A, fig. S8A**). We observed one cluster highly expressing proinflammatory cytokine *IL-1β*, as well as *CXCL8*, *CCL3*, *CXCL2* and *PTGS2*, thus termed as proinflammatory macrophages (**Fig. 5B**). Recent studies have suggested a role of IL-1β^+^proinflammatory macrophages in idiopathic pulmonary fibrosis (IPF), and in cardiac tissue remodeling(*3*, *4*). This subset also highly expresses members of the NR4A subfamily of the nuclear hormone receptor superfamily including *NR4A3, NR4A2 and NR4A1* (**Fig. 5B**), which have been identified as immediate early genes in response to various stimuli and considered as promising therapeutic targets due to their significant roles in regulating monocyte differentiation, macrophage activation and inflammatory response(*32*) and have been linked to the pathogenesis of fibrotic tissue remodeling(*33*).

**Fig. 5.**
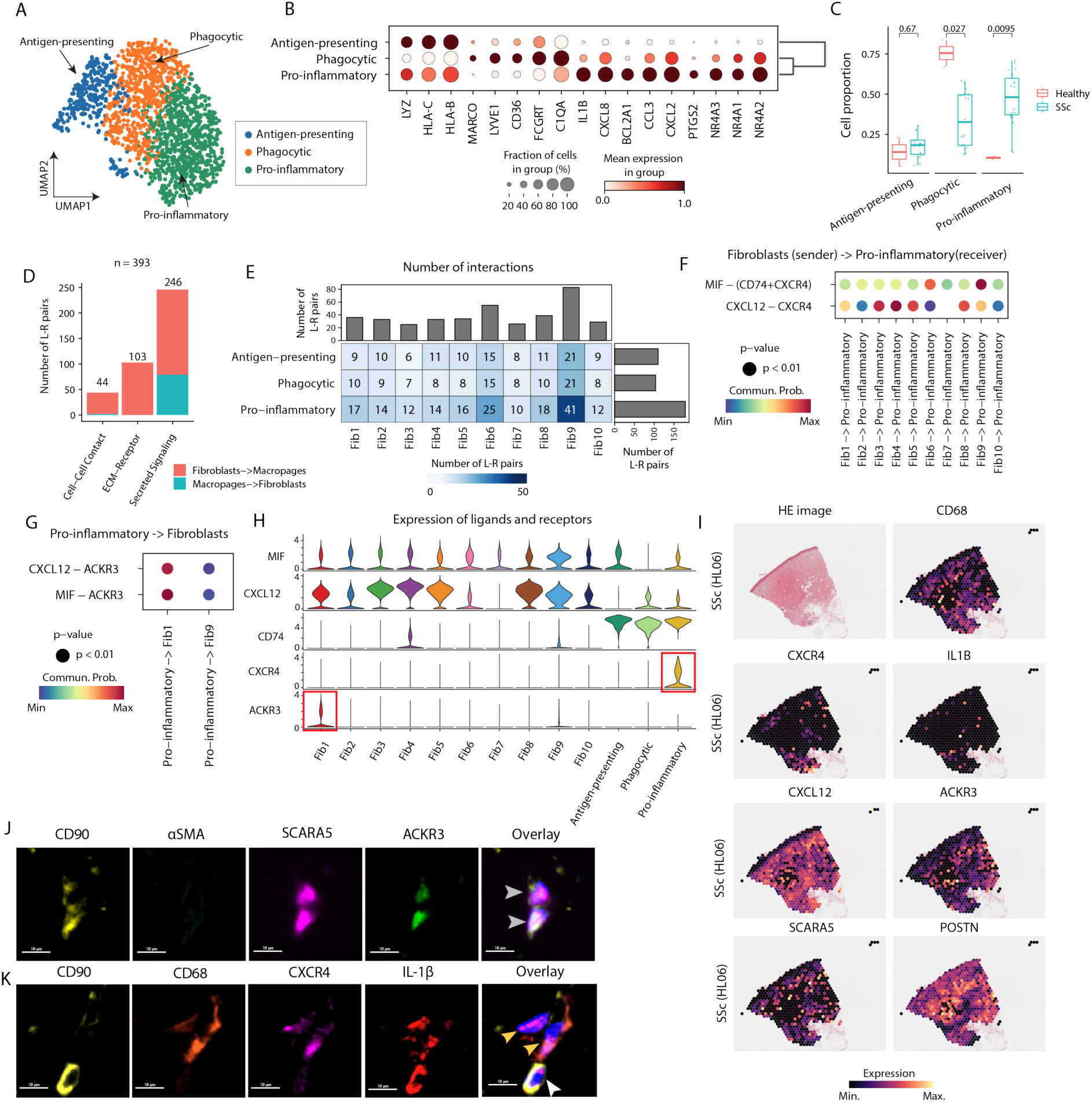
Characterization of macrophage sub-clusters. (**A**), UMAP showing sub-clusters of macrophages based on scRNA-seq data. (**B**), Dotplot showing the marker genes for each sub-cluster. (**C**), Boxplot comparing the proportion of each sub-cluster between healthy and SSc samples. Each dot represents one sample. P-values are calculated using the Wilcoxon rank-sum test (unpaired; two sides). (**D-E**), Number of significant L-R pairs between the fibroblasts and macrophages. (**F**), Heatmap showing significant ligand-receptor pairs between fibroblast subsets and proinflammatory macrophages, with fibroblast serving as signaling senders and macrophage as receivers. The colors refer to the communication probability and the sizes represent p-values. (**G**), Heatmap showing significant ligand-receptor pairs between proinflammatory macrophages and fibroblast subsets (Fib1, Fib9), as macrophage serving as signaling senders and fibroblast as receivers. (**H**), Violin plot showing the expression of ligands and receptors in scRNA-seq. (**I**), Expression of ligands and receptors in spatial space. (**J**), Representative images of TSA (Tyramide Signal Amplification) staining on the expression of ACKR3 in SCARA5^+^ fibroblast which are labeled with CD90 (indicated by grey arrows). (**K**), Representative images of TSA staining on the co-occurrence of fibroblasts (marked by CD90 and indicated by a white arrow) with IL-1β^+^macrophages, which are also positive for CXCR4 (indicated by yellow arrows). The staining in J and K were performed on adjacent tissue sections to those used for 10× Visium. Horizontal scale bars represent 10 µm.

We also identified a sub-cluster characterized by phagocytic markers, including *MARCO*, *LYVE1*, *CD36*, *FCGRT and C1QA*, therefore defined as phagocytic macrophages. The scavenging receptor CD36, known to mediate uptake in macrophage, has been implicated in the pathogenesis of tissue fibrosis(*34*, *35*). *MARCO*, a type II glycoprotein scavenger receptor that binds a broad range of anionic ligands, was found to be upregulated in tissue-resident macrophages and circulating monocytes upon activation. *MARCO*^+^ myeloid cells have been recognized as therapeutic targets for organ fibrosis in SSc(*36,37*). LYVE1-expressing macrophage has been demonstrated to form coordinated multi-cellular nest structures that are distributed proximal to vasculature(*38*). Another subset of macrophage that may be involved in the antigen presentation process is characterized by their overexpression of *HLA-B* and *HLA-C,* hereafter referred to as antigen-presenting macrophages. Indeed, the antigen-presenting macrophage and phagocytic macrophage defined here largely resembled previously identified Lyve1^lo^MHCII^hi^ and Lyve1^hi^MHCII^lo^ macrophages, which are conserved across tissues(*8*). Notably, the Lyve1^hi^MHCII^lo^ macrophages have been demonstrated to play a critical role in maintaining homeostasis, as depletion of Lyve1^hi^MHCII^lo^ macrophages exacerbated experimental lung and heart fibrosis, along with exacerbated vessel permeability, immune infiltration and collagen deposition(*8*).

The comparison of cell proportion between healthy and SSc samples showed that proinflammatory macrophages were significantly enriched in SSc (*p* = 0.0095), whereas the abundance of phagocytic macrophages was significantly decreased in SSc samples (*p* = 0.027) (**Fig. 5C**). We estimated TF activity for each cluster and observed a high activity of REL, RELA, and RELB in proinflammatory macrophages (**fib. S8B**). These TFs are key members of the NF-κB pathway that regulates immune and inflammatory responses(*39*), further supporting the functions of proinflammatory macrophages.

### Crosstalk between fibroblast and macrophage sub-clusters

Our analysis of ST data revealed a strong spatial association between macrophages and fibroblasts, underscoring their potential functional interplay within the tissue microenvironment (**Fig. 2C and Fig. 3L**). To further investigate the cellular crosstalk between specific macrophage and fibroblast subtypes, we performed a detailed cell-cell communication analysis using CellChat(*40*) in the scRNA-seq data based on ligand-receptor (L-R) interaction networks. We retained only statistically significant L-R pairs between fibroblast and macrophage sub-clusters, resulting in the identification of 393 distinct interactions. These included 246 secreted autocrine/paracrine signaling interactions, 103 ECM-receptor interactions, and 44 direct cell-cell contact-mediated interactions (**Fig. 5D**). This categorization provides a comprehensive view of the diverse modes of communication utilized by these cell types to orchestrate their roles in the tissue microenvironment. Among macrophage subsets, pro-inflammatory macrophages exhibited the highest number of interactions with fibroblast subsets (n = 179 interactions), suggesting that they are central players in fibroblast-mediated signaling (**Fig. 5E**). Particularly, Fib9 emerged as a key fibroblast subtype with the strongest connectivity to macrophages (n = 83 interactions), outpacing other fibroblast subsets in its interaction capacity.

Visualization of the specific L-R interactions revealed that macrophages receive signals from fibroblasts through multiple signaling pathways, including THBS, MIF, CXCL, LAMININ, and COLLAGEN (**fig. S8C**). Among these interactions, the receptor CXCR4 was observed to interact with fibroblast-derived ligands such as MIF and CXCL12. This interaction suggests a mechanism by which fibroblasts recruit and retain proinflammatory macrophages, thereby sustaining local inflammation and contributing to the progression of fibrosis (**Fig. 5F**). On the other hand, fibroblasts also acted as signal receivers, engaging the receptor ACKR3 to interact with MIF and CXCL12 (**fig. S8D and Fig. 5G**). Notably, *MIF* and *CXCL12* were expressed by nearly all subtypes of fibroblasts and macrophages (**Figs. 5, H and I**), underscoring their widespread involvement in cellular crosstalk.

However, the expression of *CXCR4* was specific to proinflammatory macrophages, while *ACKR3* was uniquely expressed in the Fib1 fibroblast progenitor subcluster (**Fig. 5H**). To confirm at the protein levels, we applied Tyramide Signal Amplification (TSA) high-plex immunofluorescence onto adjacent tissue sections. Our TSA staining confirmed the overlapping expression of ACKR3 in *SCARA5*^+^ fibroblast (**Fig. 5J**). Again, we observed IL-1β^+^ macrophages, which were also positive for CXCR4, closely distributed adjacent to fibroblasts (**Fig. 5K**). Together, these results revealed a highly organized and reciprocal signaling network between fibroblasts and macrophages, underscored by distinct L-R interactions. Of particular interest, the chemokine CXCL12 can bind to its classic receptor CXCR4, a G protein-coupled receptor, facilitating the recruitment of CXCR4^+^ cells, such as proinflammatory macrophages in our case. Alternatively, CXCL12 can interact with its atypical chemokine receptor ACKR3 independent of GPCR mediated signaling. ACKR3 is capable of actively internalizing its chemokine ligands CXCL12, through scavenging. This function helps to modulate the availability and gradient of CXCL12 in the extracellular environment(*41*). The selective expression of ACKR3 in fibroblast progenitors may indicate an endogenous compensation at an early stage. A dysfunction in this mechanism could lead to the subsequent CXCR4 mediated recruitment of macrophages, during the transition from fibroblast progenitor to myofibroblast over the course of SSc progression.

### A data-driven model of SSc

Taken together, we proposed a data-driven disease model of SSc based on the results of multi-omics data obtained from SSc, a prototypical systemic fibrotic condition (**Fig. 6**). An imbalance in fibroblast subsets in SSc was characterized by a decrease in *SCARA5*^+^ fibroblast progenitors and an increase in *POSTN*^+^ and *ACTA2*^+^ terminally differentiated myofibroblasts, which promote the excessive ECM deposition. The decreased phagocytic macrophages (Lyve1^hi^MHCII^lo^) may contribute to the impairment of vessel integrity. The infiltration of proinflammatory macrophages (IL-1β^+^) was observed actively interact with fibroblast subsets. These proinflammatory macrophages were overexpressed with *CXCR4*, which may mediate their recruitment by interacting with *CXCL12*. Additionally, the expression of ACKR3 on fibroblasts, particularly in progenitors under homeostatic conditions, may regulate the local levels CXCL12 by internalizing the ligand, thereby limiting excessive leukocyte recruitment and preventing unwanted tissue inflammation. However, this regulatory mechanism might diminish in effectiveness as progenitor fibroblasts decrease during the progression of SSc, ultimately resulting in uncontrolled CXCL12/CXCR4-mediated inflammatory cell attraction. This regulatory network fosters a shift from tissue homeostasis to a fibrogenic state, driving the progression of SSc. Therefore, the integration of spatial omics with scRNA-seq provides insight into the immune-stromal crosstalk involved in the pathogenesis of SSc.

**Fig. 6.**
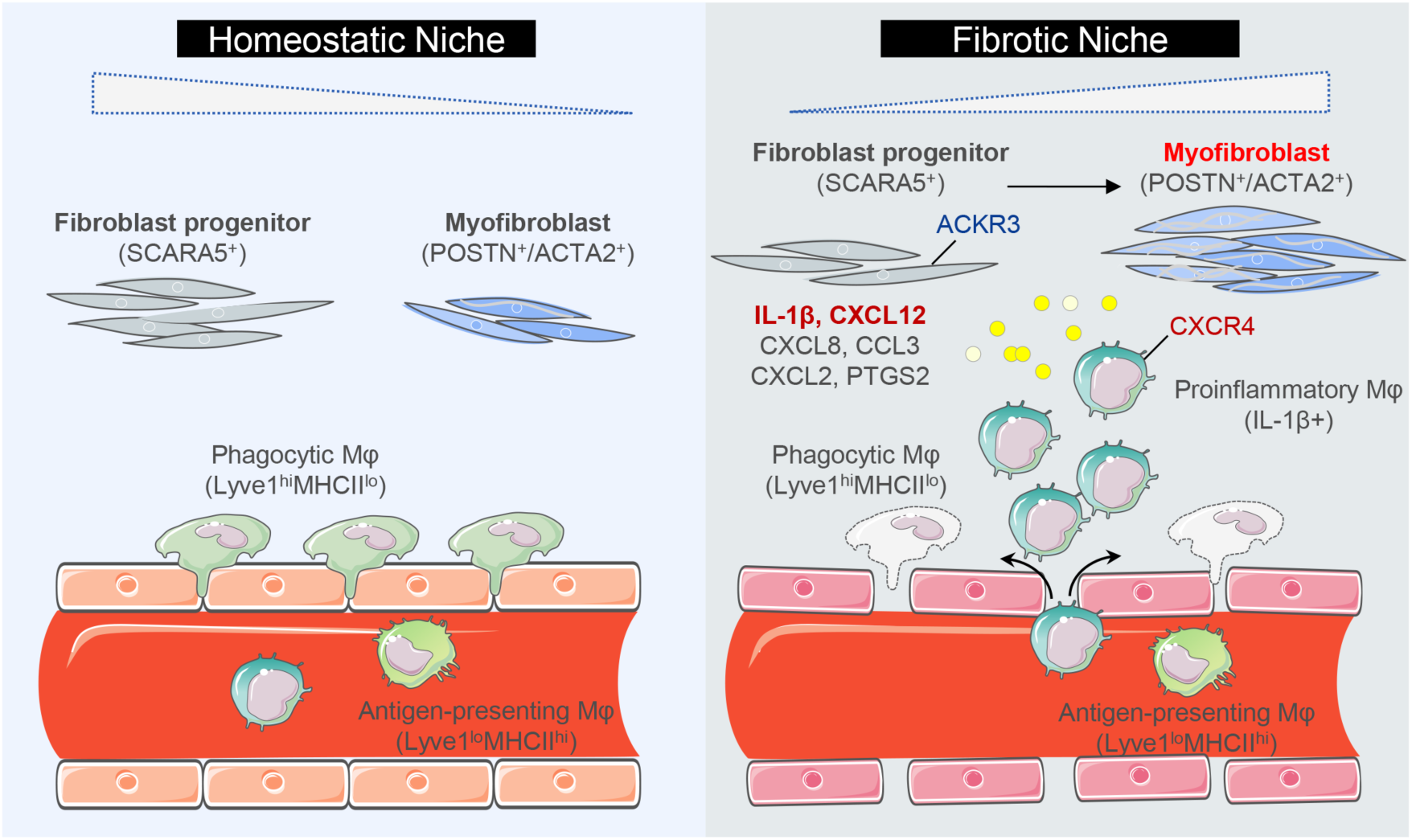
A Data-Driven Model of SSc. Schematic summary illustrates the proposed cellular interplay in SSc skin, which leads to a shift from tissue homeostasis to a fibrogenic environment. The illustration was created by using images from Servier Medical Art (http://smart.servier.com/), licensed under CC BY 3.0 (https://creativecommons.org/licenses/by/3.0/).

## DISCUSSION

SSc is a multifaceted autoimmune disease characterized by a well-known triad of manifestations: vasculopathy, autoimmunity and fibrosis. In this study, we provided a comprehensive map of the skin ecosystem in SSc using genome-wide spatial transcriptomes based on 10x Visium and Stereo-seq platforms combined with additional scRNA-seq datasets. We identified disease-associated fibroblasts and macrophages and characterized their intricate interplay within niches that potentially drive the transition from a homeostatic to a fibrogenic state. The molecular signatures of cell populations were validated through spatial proteomics, including IMC and TSA. The differentiation trajectory of fibroblasts was cross-validated using time-course bleomycin induced SSc mouse and wound-healing reindeer models. Further investigation into cell-cell communications offered potential for identifying targetable disease-driving molecules for therapeutic intervention.

We identified distinct niches enriched with fibroblasts and macrophages, showing active TGF-β signaling pathway and correlating with clinical outcomes in SSc. We then homed in on distinct cell phenotypes and spatial relationships of fibroblasts and macrophages within niches. Of note, we discovered a proportional imbalance of fibroblast subsets in SSc, with a clear reduction in SCARA5^+^ progenitor fibroblast and an upregulation in *POSTN*^+^/*ACTA2*^+^ terminally differentiated myofibroblasts compared with healthy controls. SCARA5 positive fibroblasts have been recently reported as myofibroblast progenitors in heart and kidney(*19*, *20*, *42*). A cross-tissue human fibroblast atlas, derived from 517 human samples encompassing 11 tissue types and diverse pathological states, demonstrated the substantial overlaps among *SCARA5*^+^ fibroblast and progenitor-like fibroblast, as well as between *POSTN*^+^ fibroblast and myofibroblast(*43*). Additionally, the dynamic changes of fibroblasts were not only identified through trajectory analysis in human SSc skin, but also validated in time-course models of SSc mouse and wound-healing reindeer using the publicly available datasets(*7*, *26*). These findings indicated that the differentiation from *SCARA5*^+^ fibroblast progenitors to myofibroblasts represents a conserved phenotype across different tissues and species. Targeting the shared mechanism has emerged as a strategy for basket trial design especially for rare diseases that share the same triggering or perpetuating signaling pathways, such as INBUILD study(*44*).

Importantly, our data revealed the baseline gene expression ratios of marker genes for myofibroblast versus fibroblast progenitor indicate progression in skin fibrosis and an incomplete response to standard treatment, assessed using a composite responsiveness criteria(*29*) based on clinically significant changes at 52-week follow-up. Pretreatment stratification of SSc patients using expression ratios revealing dynamic changes in fibroblast states underscores its vital role in precision medicine for predicting treatment response likelihood. Furthermore, future studies are necessary to elucidate the spatiotemporal changes occurring within fibrogeneis microenvironment by examining longitudinal biopsies throughout the trajectory of the disease and during treatment.

Of note, we revealed an intense interaction between IL-1β^+^ proinflammatory macrophages and fibroblast subsets in SSc. Infiltrating macrophages activate fibroblasts by secreting IL-1β, whilst systemic IL-1β inhibition or targeted IL-1β deletion in macrophages prevents fibroblast activation(*3*, *4*, *45*). Furthermore, we found this crosstalk was predicted to be mediated by the interaction of CXCL12 with CXCR4 in macrophages and ACKR3 in fibroblast progenitors. Indeed, we also found the deregulation of Lyve1^hi^MHCII^lo^ macrophages in SSc, which are known to play a protective role by maintaining vascular integrity and preventing immune cell infiltration in fibrotic conditions(*8*). These sequential events such as tissue entry and immune infiltration occuring in SSc may shift the hemostatic condition towards a profibrotic environment. Importantly, the inability to constrain CXCL12 overexpression in the milieu through internalization by fibroblast progenitors via ACKR3 leads to CXCR4-mediated proinflammatory macrophage recruitment, which is reflective of the fine-tuned regulation by local fibroblasts.

Our research aimed to explore the potential to halt the disease progression by interrupting the crosstalk between pathological cells. Spatial transcriptomics provides a unique advantage in examining the structured compartment and the cellular composition and states within specific microenvironments. However, the 10x Visium platform does hold significant limitations, such as the low resolution of 55-um per spot. To overcome this limitation and increase granularity, we employed a higher-resolution spatial transcriptomics technology, FFPE-based Stereo-seq, achieving a resolution of 0.5 um per spot. To further validate the genome-wide spatial data, we applied additionally spatial proteomics, including IMC and TSA, which offering the molecular expression and distribution of the identified cells in SSc skin at single-cell level with the resolutions of 1 um^2^ and 0.25 um^2^ per pixel, respectively. To achieve a comprehensive description of the skin landscape in SSc, we also recognize that three-dimensional profiling of tissue space and subcellular localization of protein signals will significantly expand our understanding in SSc pathogenesis.

In summary, spatial multi-omics combined with scRNA-seq helps to elucidate the entanglement of fibroblasts and macrophages, forming a connectivity network in the pathogenesis of SSc. Within this network, fibroblasts and macrophages with distinct phenotypes contribute to tissue fibrotic remodeling in a coordinated manner. We thus used this strategy to facilitate target discovery, identifying potential intervention axes for fibrosis.

## MATERIALS AND METHODS

### Study Design

The objective of this study was to unbiasedly characterize the cellular niche and neighbourhood interaction driving SSc fibrosis, providing spatiotemporal insights into potential target identification and precise stratification. We thus obtained skin punch biopsy samples from patients with patients with early and diffuse cutaneous involvement of SSc, and as well as from healthy controls. We generated spatially resolved transcriptome maps using two complementary spatial technologies: 10x Visium and Stereo-seq (spatial enhanced resolution omics sequencing). By integrating these datasets with publicly available scRNA-seq datasets, we constructed a population-scale human skin transcriptome atlas, which enabled us to study the cellular and molecular landscape of SSc with spatial resolution. We then conducted the subsequent analyses, including cell-type deconvolution, tissue structure identification, disease-associated subclustering, trajectory analysis, spatial colocalization, and ligand-receptor analysis. The findings were validated using spatial proteomic techniques such as imaging mass cytometry (IMC) and tyramide signal amplification (TSA), as well as time-course scRNA-seq data of SSc mouse and wound-healing reindeer models. Furthermore, we analyzed the associations of these findings with clinical outcomes and treatment responsiveness in SSc (**Fig. 1A**).

**Further details on Materials and Methods can be found in the Supplementary Materials.**

## List of Supplementary Materials

Supplementary Materials and Methods

Fig. S1: Quality check of spatial transcriptomics data.

Fig. S2: Analysis of publicly available scRNA-seq data.

Fig. S3: Characterization of cell-type niches in human skin.

Fig. S4: Validation of cellular neighbourhood (CN) by imaging mass cytometry (IMC) in healthy and SSc skin.

Fig. S5: Sub-clustering of fibroblasts.

Fig. S6: Trajectory analysis of fibroblast differentiation.

Fig. S7: Cross-Species Validation of Fibroblast Dynamics.

Fig. S8: Sub-clustering of macrophages.

Table. S1: Clinical characterization of the patients with SSc undergoing skin biopsy for spatial transcriptomic study.

Table. S2: Quality statistics for 10x Visium and Stereo-seq data.

## Acknowledgments

Natural Science Foundation of China (NSFC) 82371818 (ML)

Joint Sino-German research project from NSFC and Deutsche Forschungsgemeinschaft (DFG, German Research Foundation) 82161138022 (JHWD and HZ)

Huashan Cultivation research 30302164001 (ML)

Huashan Innovation project 2024CX01 (ML)

Shanghai Oriental Youth Talent program DFYCQN07 (ML)

Rappaport MGH Research Scholar Award (LP)

## Acknowledgments

We would particularly like to acknowledge my colleagues, Division of Rheumatology of Huashan Hospital, Fudan University, for their wonderful collaboration and support. We are also deeply grateful to the patients who generously provided the samples used in this study.

Their contributions are invaluable to our research.

## Author contributions

Conceptualization: ML and ZL

Methodology: ZL, ML, ARR, WX, LH, XS, YNL, AEM, WY

Investigation: ZL, ML, ARR

Visualization: ZL, ML, ARR

Funding acquisition: ML, JHWD, LP and HZ

Project administration: ML

Supervision: ML, ZL, JHWD, RH

Writing – original draft: ML and ZL

Writing – review & editing: ZL, ARR, WX, LH, XS, YNL, AEM, WY, HZ, LP, JHWD, RH, ML

## Data and materials availability

The spatial transcriptomics data presented in this study are available at the Zenodo data archive (https://zenodo.org/records/14577696). The scRNA-seq data for human skin were downloaded from GEO with the accession number <u>GSE195452</u> (Gur2022) and <u>GSE138669</u> (Tabib2021). A cross-tissue human fibroblast atlas was analyzed with the interface website http://pan-fib.cancer-pku.cn (Gao2024). The scRNA-seq data of full-thickness injuries in skin from reindeer was analyzed with the interface website www.biernaskielab.ca/reindeer_atlas (Sinha2022). The scRNA-seq data for mouse skin were downloaded from the Zenodo data archive (https://zenodo.org/records/7455989). The codes used in this study are available on Github: https://github.com/lzj1769/Spatial_SSc_project

## Abbreviations

CRP: C-reactive protein
mRSS: Modified Rodnan Skin Score
PLT: platelet count
C3: complement component 3
ILD: interstitial lung disease
ALT: alanine aminotransferase
C4: complement component 4
ESR: erythrocyte sedimentation rate
HGB: hemoglobin
LDH: lactate dehydrogenase
AST: aspartate aminotransferase
CK: creatine kinase
DU: digital ulcer
IgG: immunoglobulin G
sPAP: systolic pulmonary arterial pressure
SpO2: oxygen saturation
Cr: creatinine
TnT: troponin T
EUSTAR activity index: European Scleroderma Trials and Research Group activity index
r-GT: gamma-glutamyl transferase
TB: total bilirubin
WBC: white blood cell count
Non-RP Disease Course: non-raynaud’s phenomenon disease course).

## Supplementary Materials and Methods

### Human skin biopsy collection and processing

Skin biopsies from 14 SSc subjects and 4 age- and gender-matched healthy controls were obtained and proceeded for spatial transcriptomics using the 10x Visium platform. All skin biopsies of SSc patients were taken on the forearm 15 ± 2 cm proximal from the styloid process of the ulna using disposable biopsy punches (pfm medical, #48301, Gifu, Japan). The epidemiological and clinical features of the SSc patients are presented in table S1. After excluding 2 low-quality SSc samples (**fig S1**), we finally included 14 skin biopsies in total, from10 SSc patients and 4 healthy controls. SSc was diagnosed according to the American College of Rheumatology (ACR) /European Alliance of Associations for Rheumatology (EULAR) 2013 criteria(*46*). In particular, we enrolled SSc patients who had diffused cutaneous involvement and had experienced their initial non-Raynaud’s phenomenon symptoms for less than three years. Among the SSc patients, 40% were female and 60% were male. The median age at biopsy of SSc patients was 56.0 years (range 27-73). The median disease duration from the first non-RP symptoms was 8 months (range 4-48). The median mRSS was 21 (range 8-41). 40% of SSc patients suffered from interstitial lung disease (ILD). All the SSc patients included in our study were treatment-naive and received the standard treatment of MMF for consecutive 52 weeks following skin biopsy. The responsiveness was defined according to the criteria of the revised ACR-CRISS (American College of Rheumatology Composite Response Index in Systemic Sclerosis), which is the improvement in ≥ 3 of 5 core clinical measurements with Minimal Clinically Important Difference (MCID) of 25% for modified Rodnan skin score (mRSS), 5% for percent predicted forced vital capacity (FVC%), 20% for health assessment questionnaire-disability index (HAQ-DI), 20% for patient (PGA) and clinician (CGA) global assessments(29). SSc patients with progression of skin fibrosis, which is defined as increase in modified Rodnan skin score (mRSS) ≥ 25% from baseline in the 52-week follow-up visit after baseline. All human studies were approved by the ethical committees of Huashan Hospital of Fudan University. All patients and controls signed a consent form approved by the local institutional review board after informed consent.

### Spatially profiling human skin with 10x Visium

Formalin-fixed paraffin-embedded (FFPE) skin blocks from a total of 18 subjects were sectioned at 5 um thickness. These sections were transferred to match the 6.5mm^2^ oligo-barcoded capture areas on the 10x Visium platform (FFPE v2) according to the manufacturer’s protocol. Prior to this, archived tissues from paraffin-embedded blocks were tested for RNA quality control (DV200 > 30%), before being mounted on the 10× CytAssist (6.5 mm × 6.5 mm) for library preparation and sequencing (∼50,000 reads/spot). Hematoxylin and eosin (H&E) staining was conducted for morphological assessment. The Visium Human Transcriptome Probe Set v2.0 was applied. The final library pool was sequenced on the Illumina Novaseq 6000 instrument using 150-base-pair paired-end reads.

### Spatially profiling human skin with Stereo-seq

To achieve an enhanced resolution and whole transcriptome in-situ analysis of skin samples, we employed Stereo-seq technology according to the previous protocols(*16*, *47*), which enables the spatial detection of transcriptome-wide gene expression at a resolution of 0.5 um. Briefly, 5-um FFPE skin sections from 3 SSc patients and 1 healthy control, which are overlapping with the ones who underwent the examination of 10x Visium, were subjected to adhere to the Stereo-seq chips which have DNA nanoball (DNB) bins with 220 nm diameter and a center-to-center distance of 500 nm. After methanol fixation, the formal sections were preceded with ssDNA staining and the adjacent sections with H&E staining for morphological assessment. Next, permeabilization of the tissue was performed to release mRNA. cDNA products are synthesized by reversed transcription. After amplification, DNBs were then generated via rolling cycle amplification (RCA) and loaded onto the patterned chips (65mm×65mm). Next, to determine the distinct DNB-CID sequences at each spatial location, paired-end sequencing was performed on a DNBSEQ-T7 sequencer (MGI Research, Shenzhen, China) with a PE75 sequencing strategy.

### Processing 10x Visium and Stereo-seq data

For 10x Visium data, we used SpaceRanger software (v2.0.0) to pre-process the raw sequencing data based on the reference genome hg38. For each sample, we filtered the spots by number of detected genes (< 100). For Stereo-seq data, we leveraged nucleic acid staining from the same section to segment cells by projecting the staining image to the Stereo-seq chips. The watershed algorithm was applied to the staining image for single cell segmentations. For each of the segmented cells, UMI from all DNB within the corresponding segmentation were aggregated per-gene and then summed to generate a cell by gene matrix for downstream analysis. To overcome the low RNA capture efficiencies on single DNB spots at a resolution of 500 nm, the raw spatial expression matrix was convoluted into larger pseudo-spots with a 50×50 window size (bin50 for short), or more precisely, as 25 µm squares. We then filtered the spots by number of detected genes (< 50) and mitochondrial genes (> 10%).

### Analysis of publicly available scRNA-seq data

To build an atlas of human skin cells, we downloaded scRNA-seq data from Gene Expression Omnibus (GEO) with accession numbers GSE195452 (denoted by Gur2022)(*10*) and GSE138669 (denoted by Tabib2021)(*9*). For the Gur2022 dataset, we only kept the cells from the skin. We further filtered the cells by number of UMIs (< 500), number of detected genes (< 300 or > 5000), and mitochondrial genes (> 30%). We also filtered the genes by the number of cells (n = 100). We re-annotated the cells by merging the 49 original labels to 16 major cell types as follows: Fibroblast (Fibro_ATAC2, Fibro_Bad, Fibro_COCH, Fibro_COMP, Fibro_IGFBP2, Fibro_LGR5, Fibro_MYOC1, Fibro_MYOC2, Fibro_POSTN, Fibro_PTGDS, Fibro_POSTN_PTGDS), T cells (sT_Effector, T_Effector, T_GD, T_Naive, sTreg_CXCR4, sTreg, Treg, sT), NK cells (NK, NK_XCL1, NK_XCL1_CXCR4), cDC (DC, DC_CCL22, DC_CXCL10, DC_XCR1), Macrophages (Mf, Mf_TREM2), Monocytes (Mo_CD16, M_CD16_IL1B, M_IL1B, Mo), Mast cells (bMast_CLC, Mast), B cells (B, B_CXCR4), Pericytes (Peri_RGS5, Peri_TGFBI), Endothelial cells (Vascular_ACKR1, Vascular_RBP7), Keratinocytes (KRT1_KRT10, KRT14_ACTA2, KRT14_S100A2_GJA1). For the Tabib2021 dataset, we filtered the cells by UMIs (<500), number of detected genes (<300 and >5000), and mitochondrial gene expression (>5%). Since the original annotation was unavailable, we clustered and re-annotated the cells. This recovered 13 major cell types, among which 10 cell types were also identified by the Gur2022 dataset. To integrate these two datasets, we merged the count matrices and normalized the data using Scanpy(*48*). Next, we used the Harmony algorithm(*49*) to correct the batch effects between the datasets and visualize the integrated cells by UMAP.

### Cell type deconvolution of 10x Visium data

We used cell2location(*50*) and the above scRNA-seq data as a reference to estimate cell type composition for each spot in the spatial transcriptomics samples. Specifically, we first identified the signatures of each cell type using the scRNA-seq data based on a Negative binomial regression model after accounting for the batch effects from different datasets. Next, we removed all mitochondrial genes, subsetting the reference data and spatial data using the identified signatures. We ran the function Cell2location (N_cells_per_location=5; detection_alpha=20) to estimate posterior distributions of cell abundance. We normalized the estimated cell type composition for each spot to [0, 1], which was used for visualization and downstream analysis.

### Inference of pathway activity for 10x Visium data

We used decoupeR(*51*) (v1.8.0) to infer the pathway and transcription factor activity. Specifically, for each sample, we normalized the expression counts matrix and identified highly variable genes using Scanpy(*48*). Next, we obtained a collection of pathways and their target genes using the function get_progeny (organism=’human’, top=500). The pathway enrichment scores were inferred using the multivariate linear model by running the function run_mlm.

### Integration of scRNA-seq and Stereo-seq data

We used Seurat (v5.1.0) to integrate the scRNA-seq and Stereo-seq data. Specifically, we merged the Stereo-seq samples and then identified a set of anchors between these two datasets using the function FindTransferAnchors (reduction = “cca”) based on the shared genes (n = 11110). Next, we transferred the cell type labels from scRNA-seq to Stereo-seq data using the function TransferData (weight.reduction = “cca”, dims = 1:30). To integrate the data in a common embedding space, we also transferred the batch-corrected PCA components (n = 30) from scRNA-seq to Stereo-seq data with the function TransferData. Next, we run the function RunUMAP to generate a co-embedding space to visualize integrated scRNA-seq and Stereo-seq data.

### Identification of cell-type composition niches in 10x Visium data

We combined the cell type proportions estimated by cell2location from all samples to. Next, we ran the function neighbors (n_neighbors=30) to create a neighborhood graph of all spatial spots based on the cell type proportions. An UMAP embedding was generated using the function umap with default settings in Scanpy. We clustered the spots with the function leiden (resolution = 0.5), identifying 12 niches. To annotate the niches, we computed the average proportion for each cell type within each niche. We next normalized the values by z-score transformation and clustered the niches using hierarchical approach.

### Sub-clustering of fibroblasts

We subsetted the fibroblasts and removed the samples with less than 100 cells to allow for robust data integration. Next, we clustered the cells with the Leiden algorithm(*52*) (resolution =1) based on the harmonized principal components. We removed one population with high mitochondrial genes, resulting in a total of ten clusters identified. The marker genes for each cluster were identified using the function rank_genes_groups (method=’wilcoxon’) from Scanpy. To map these sub-clusters to spatial space, we selected the top 100 markers for each cluster and then computed module scores using the function AddModuleScore from Seurat(*53*).

### Analysis of scRNA-seq data from mouse skin

We downloaded scRNA-seq data from the Zenodo data archive (https://zenodo.org/records/7455989). Cells were filtered by the number of UMIs (< 2000 or > 40000), number of detected genes (< 200 or > 6000) and fraction of mitochondrial genes (> 5%), retaining 18503 cells. Genes were filtered by the number of cells (< 50). Next, we normalized the count matrix using the functions normalize_total and log1p from Scanpy. Highly variable genes were selected by the function highly_variable_genes. We clustered the cells with the Leiden algorithm (resolution = 0.5) based on the 30 principal components and identified 25 populations which were further grouped into 14 cell types based on their canonical markers. We next subsetted all fibroblasts and clustered the cells using the Leiden algorithm (resolution=0.5), obtaining 10 sub-clusters. We noticed one cluster showing a low number of UMIs and detected genes, presumably corresponding to low-quality cells. We therefore removed this cluster and rename the remaining clusters as Fib1-9.

### Analysis of scRNA-seq data from reindeer

The scRNA-seq data of full-thickness injuries in skin from reindeer was analyzed with the interface website (www.biernaskielab.ca/reindeer_atlas)(7).

### Trajectory analysis of fibroblasts

We used ArchR(*54*) to perform trajectory analysis. Specifically, we inferred two pseudo-time trajectories using the function AddTrajectory to characterize the differentiation processed from Fib1 to Fib2 (denoted by trajectory 1), and from Fib1 to Fib9 (denoted trajectory 2) based on the principal components (n = 30). The trajectory 1 involved six subsets (Fib1, Fib3, Fib5, Fib4, Fib10, and Fib2), and the trajectory 2 included only Fib1 and Fib9. The trajectories were visualized by the function TrajectoryPlot. To identify associated genes for each trajectory, we obtained the expression matrix along the trajectory using the function GetTrajectory (smoothWindow = 11). Next, we visualized the dynamics of gene expression with the function plotTrajectoryHeatmap and selected the top 10% highly variable genes across the trajectory.

### Spatial relationship analysis between fibroblast subsets and other cell types

We used two approaches to infer spatial relationships between the subsets of fibroblasts and other major cell types. The first approach was based on Spearman correlation between the module score of each subset and the proportion of each major cell type across all spatial spots. This was performed for each 10x Visium sample independently, and results were averaged across all samples. The second approach was based on Misty. Specifically, we usd Misty to build a predictive model using the cell type proportions as features and module scores as target. This allows us to learn the spatial interactions between spots.

### Sub-clustering of macrophages

We subsetted all macrophages from the integrated scRNA-seq atlas and removed samples with less than 50 cells. Next, clustered the cells using the Leiden algorithm(*52*) (resolution = 0.3) based on the harmonized principal components, identifying three sub-clusters. The marker genes for each cluster were identified using the function rank_genes_groups (method=’wilcoxon’) from Scanpy. To map these sub-clusters to spatial space, we selected the top 100 markers for each cluster and then computed module scores using the function AddModuleScore from Seurat(*53*).

### Cell-cell communication analysis between fibroblasts and macrophages

We merged the fibroblast and macrophage subclusters to create a Seurat object. Next, we normalized the count matrix using the function NormalizeData from Seurat and identified highly variable genes using the function FindVariableFeatures. We created a CellChat object and grouped the cells based on the sub-clustering results as described above. We selected the database CellChatDB.human and excluded ligand-receptor pairs from non-protein signaling. The cell-cell communication network was inferred using the function computeCommunProb (type = “triMean”) and was filtered using the function filterCommunication (min.cells = 10).

### Imaging mass cytometry (IMC)

We performed data mining using our published IMC dataset(*15*), which included FFPE skin biopsy sections from 19 SSc patients and 14 matched controls. The IMC panel included up to 38 lanthanide labeled antibodies as well as Iridium 193 and 191 isotopes, which were designed to target epitopes specific for different subsets of endothelial cells, endothelial precursor cells, immune cells and mesenchymal cells, as well as structural markers used for tissue segmentation. Data acquisition was performed on a Helios time-of-flight mass cytometer coupled to a Hyperion Imaging System (Standard Biotools). After single-cell segmentation and data transformation and normalization, we obtained the single-cell dataset with spatial information alongside. We then proceeded with dimensionality reduction, unsupervised clustering, and subsequent cell type annotation. To investigate the local microenvironment of individual fibroblast subpopulations, we identified cellular neighborhoods (CNs) by k-means clustering on the composition of neighboring cells as described(*14*).

### Tyramide signal amplification (TSA)

FFPE skin tissue sections obtained from 4 SSc patients and 4 control donors were stained using a TSA system as previously described(*15*, *55*). Briefly, the sections were proceeded with dewaxing, rehydration, and antigen retrieval with Tris-EDTA(pH 9.0) for 15 minutes. Endogenous peroxidase activity was neutralized with 3% H_2_O_2_ for 10 minutes, followed by a 30-minute blocking step with antibody diluent/block buffer (ARD1001EA, AKOYA). Next, sections were incubated for 90 min at room temperature with the following primary antibodies of the indicated dilutions: SCARA5 (ab118894, 1:200, Abcam), CD90 (ab226123, 1:200, Abcam), ɑ-SMA (#19245, 1:200, Cell signaling), CD68 (ab192847, 1:200, Abcam), IL-1β (ab283818, 1:200, Abcam), CXCR4 (ab181020, 1:200, Abcam) and ACKR3 (20423-1-AP, 1:200, Proteintech). After multiple rounds of signal amplification with Opal fluorophores and subsequent antibody stripping, sections were stained with DAPI. Slides were scanned using a PhenoImager Fusion scanner (Akoya Biosciences) to distinguish the spectra of the different fluorescence channels. All images were captured and presented using the same equipment setting. Merging channels was conducted using ImageJ / Fiji software (Release 2.16.0, https://fiji.sc).

### Statistics

Quantitative data are shown either as violin plots, bar graphs or scatter plots. Data in bar graphs are presented as median ± interquartile range (IQR) with individual data points plotted as dots. Statistical significance was calculated using R (4.4.2). Further statistical analysis on clinical correlation was also performed using GraphPad prism 8 version 8.3.0. We used nonparametric tests in this study. Two-sided Wilcoxon rank sum tests were used to perform two-group comparisons in Fig. 2D, Fig. 3C, Fig. 4G, Fig. 4H, Fig. 5C, and figs. 5 (C and D). Non-parametric Spearman correlation was used to assess relationships between continuous variables in Fig. 2I, Fig. 3H, Figs. 4 (C and E), and figs. S5 (C and D). *P*-values < 0.05 were considered statistically significant.

## SUPPLEMENTAL FIGURES

**Fig. S1.**
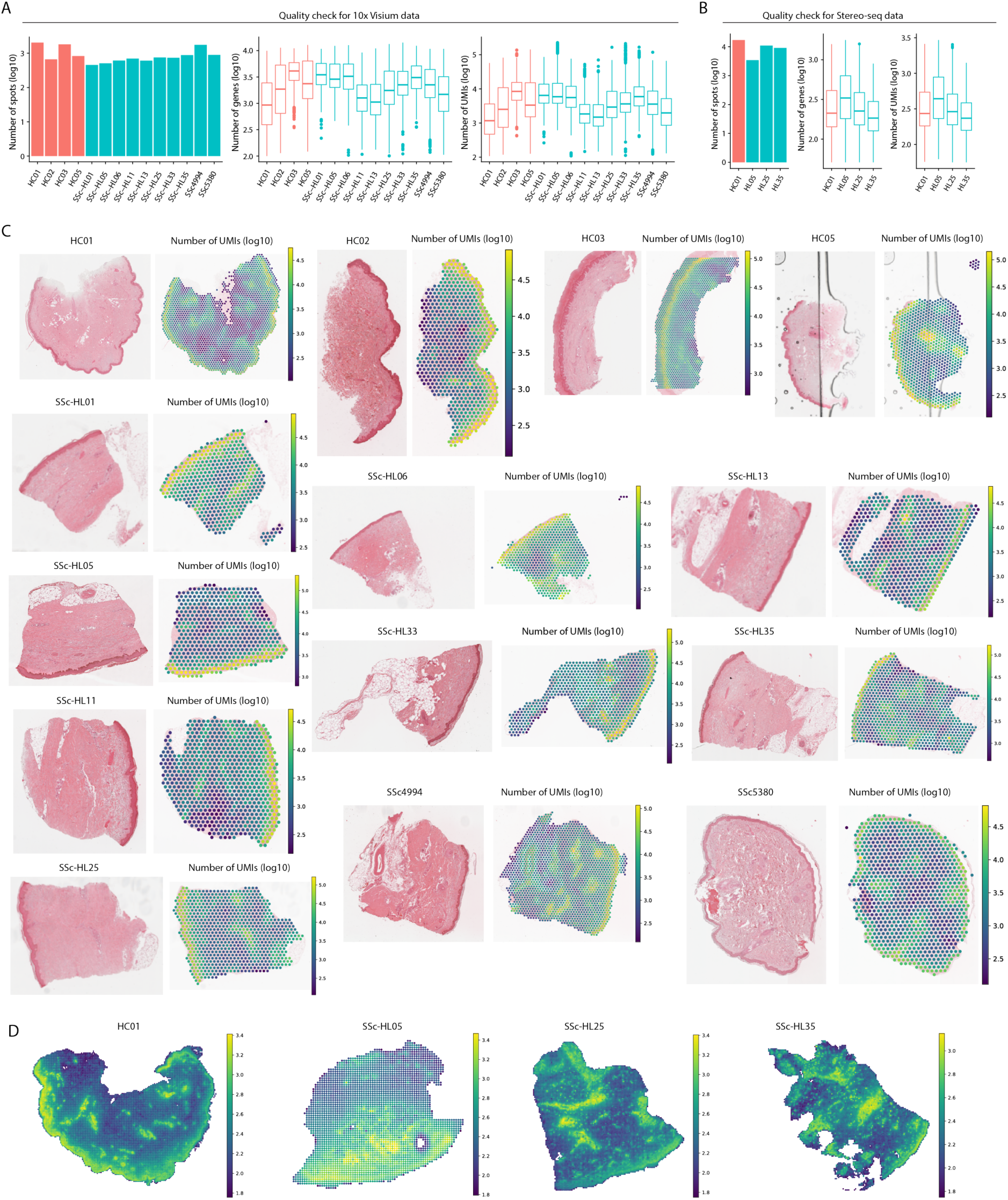
Quality check of spatial transcriptomics data. (**A**), Left: number of detected spots per sample in 10x Visium data. Middle: number of genes per spot in each sample. Right: number of UMIs in each sample. (**B**), Left: number of detected spots per sample in Stereo-seq. Middle: number of genes per sample. Right: number of UMIs in each sample. (**C**), Visualization of H&E image and number of UMIs in spatial space for 10x Visium data. (**D**), Visualization of number of UMIs in spatial space for Stereo-seq data.

**Fig. S2.**
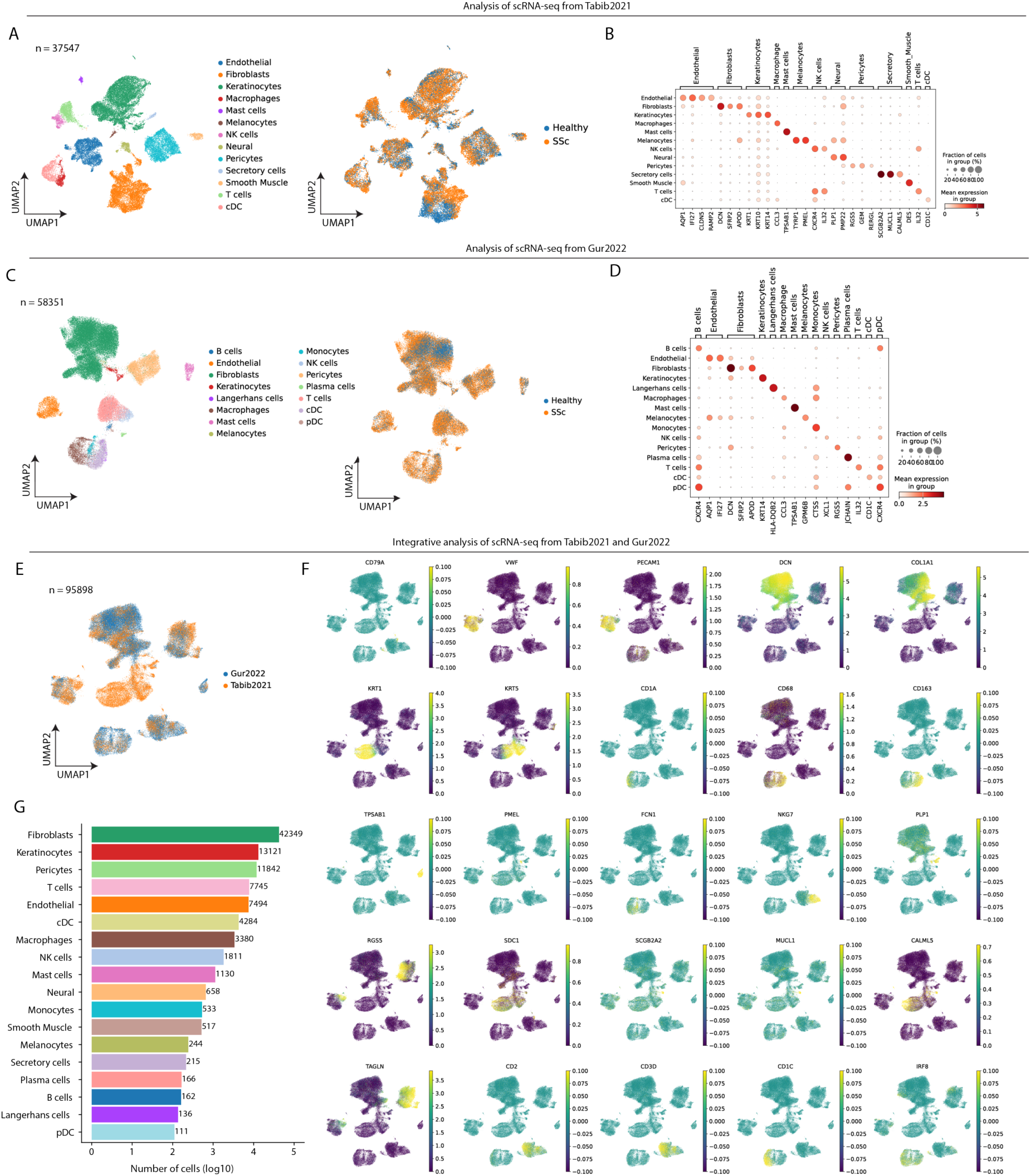
Analysis of publicly available scRNA-seq data. (**A**), UMAP showing scRNA-seq from the Tabib2021 dataset as colored by cell types and conditions. Heatmap showing cell-type-specific marker genes. (**B**), UMAP showing scRNA-seq as colored by cell types and conditions. (**C**), Same as A for the Gur2022 dataset. (**D**), Same as B for the Gur2022 dataset. (**E**), UMAP showing the integrated scRNA-seq data colored by different datasets. (**F**), Visualization of cell-type-specific marker genes. (**G**), Barplot showing number of cells for each cell type.

**Fig. S3.**
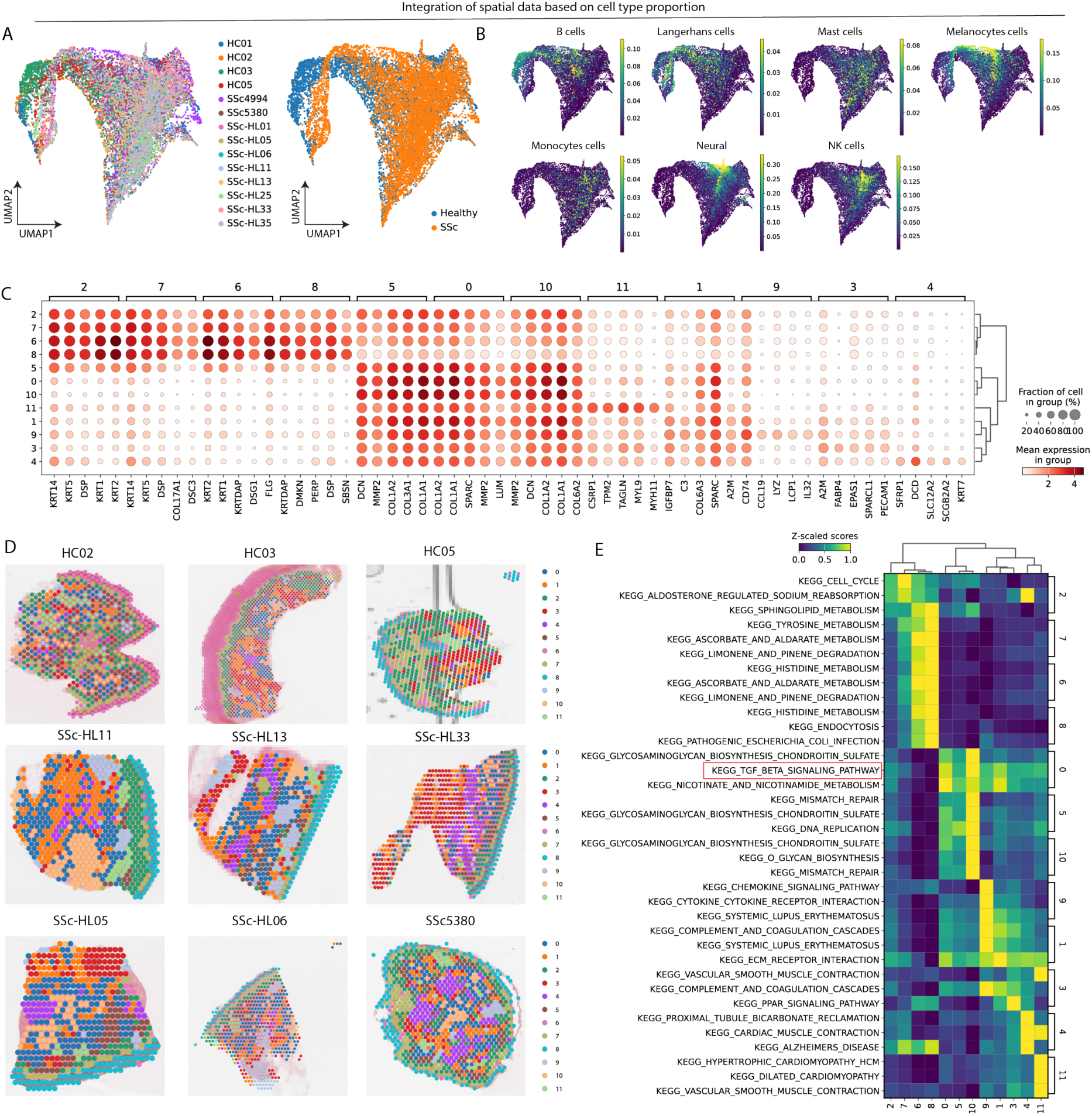
Characterization of cell-type niches in human skin. (**A-B**), UMAP embedding of the integrated spatial data as colored by samples, conditions, and the fraction of each cell type. (**C**), Marker genes of each cell-type niche. (**D**), Visualization of cell-type niches in spatial space. (**E**), KEGG pathway enrichment analysis for each cell-type niche.

**Fig. S4.**
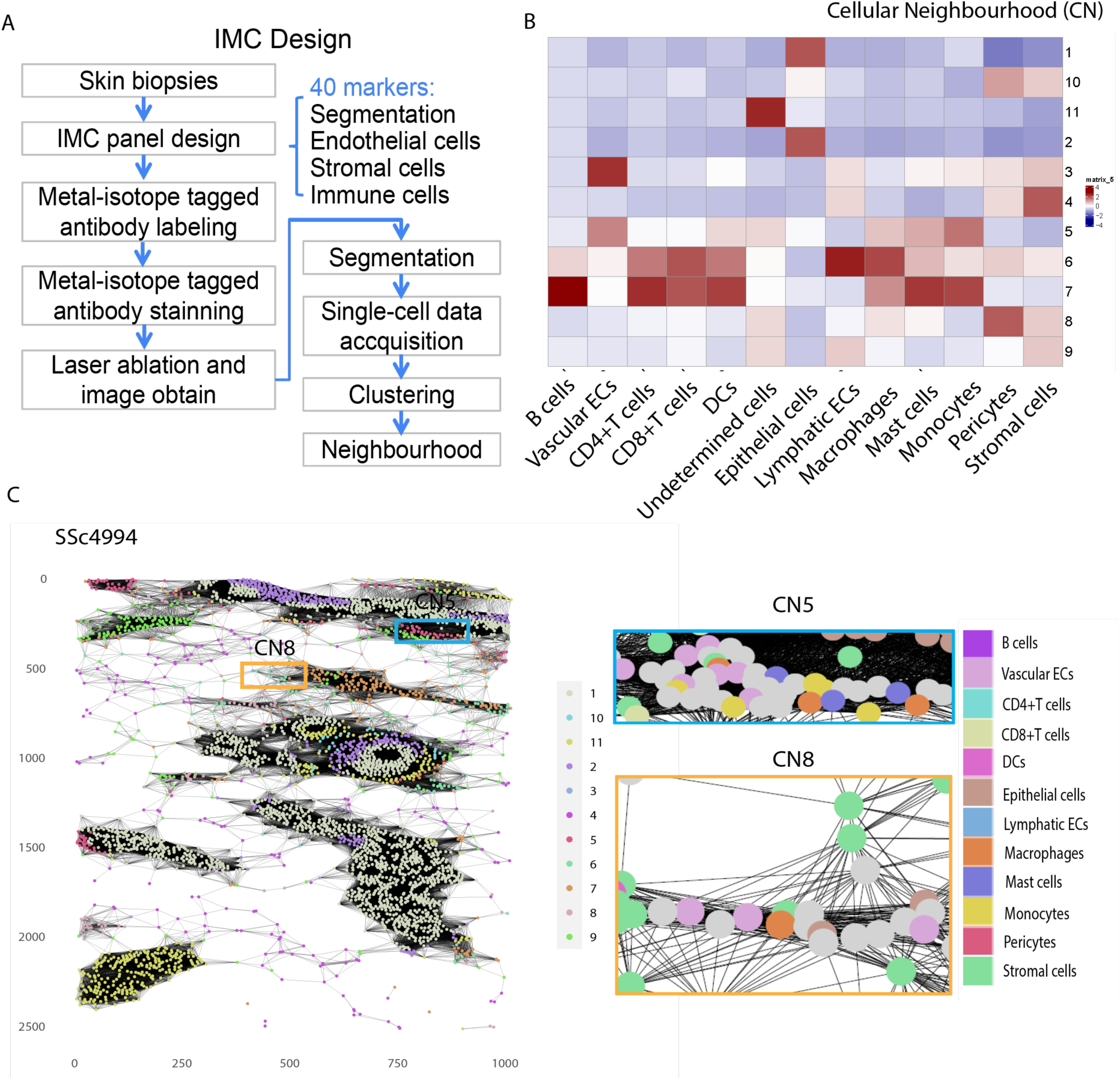
Validation of cellular neighborhood (CN) by imaging mass cytometry (IMC) in healthy and SSc skin. (**A**), Study design of IMC, as a validation spatial approach. We collected skin biopsies from 19 SSc patients and 14 healthy controls as we previously described(*15*). (**B**), Heatmap showing the enrichment of cell types in CNs. Each column represents a cell type, and each row represents a CN. Colors refer to the proportions of each cell type across different CNs. (**C**), Left: CN community detected using IMC. Colors refer to cellular clusters. Right: CN5 and CN8 are highlighted and colored by cell types.

**Fig. S5.**
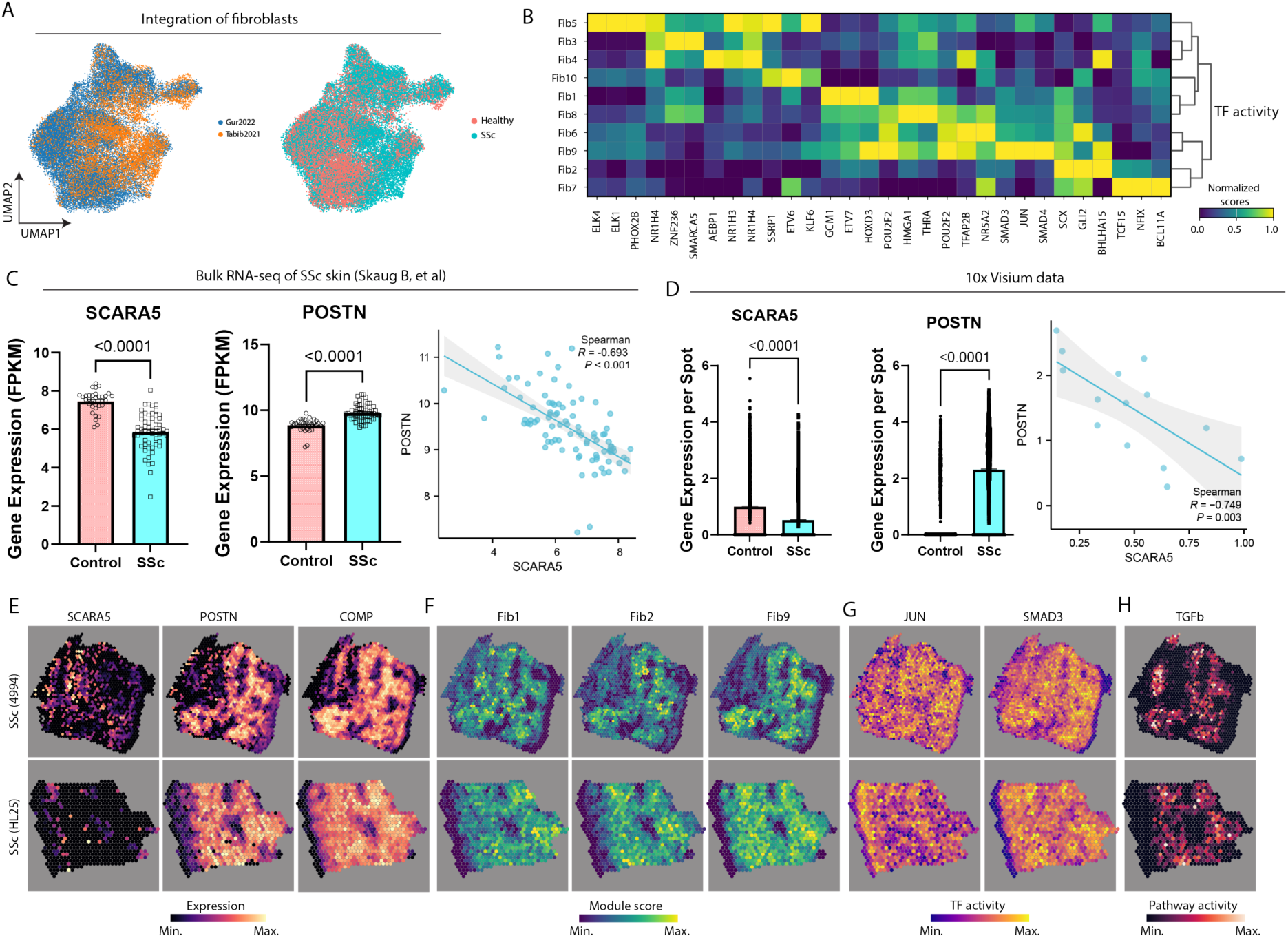
Sub-clustering of fibroblasts. (**A**), UMAP embedding of the integrated scRNA-seq fibroblasts as colored by datasets and conditions. (**B**), Heatmap of top four marker transcription factors (TFs) for each sub-cluster-based TF activity score. (**C**), Left: barplot comparing the expression of *SCARA5* and *POSTN* between healthy and SSc conditions using bulk RNA-seq. Right: scatter plot showing the Spearman correlation between *SCARA5* and *POSTN* across patients. (**D**), Left: barplot comparing the expression of *SCARA5* and *POSTN* between healthy and SSc conditions using Visium data. Right: scatter plot showing the Spearman correlation between *SCARA5* and *POSTN* across Visium samples. (**E-H**), Visualization of marker genes (E), module scores (F), TF activity (G) and pathway activity (H) in spatial space.

**Fig. S6.**
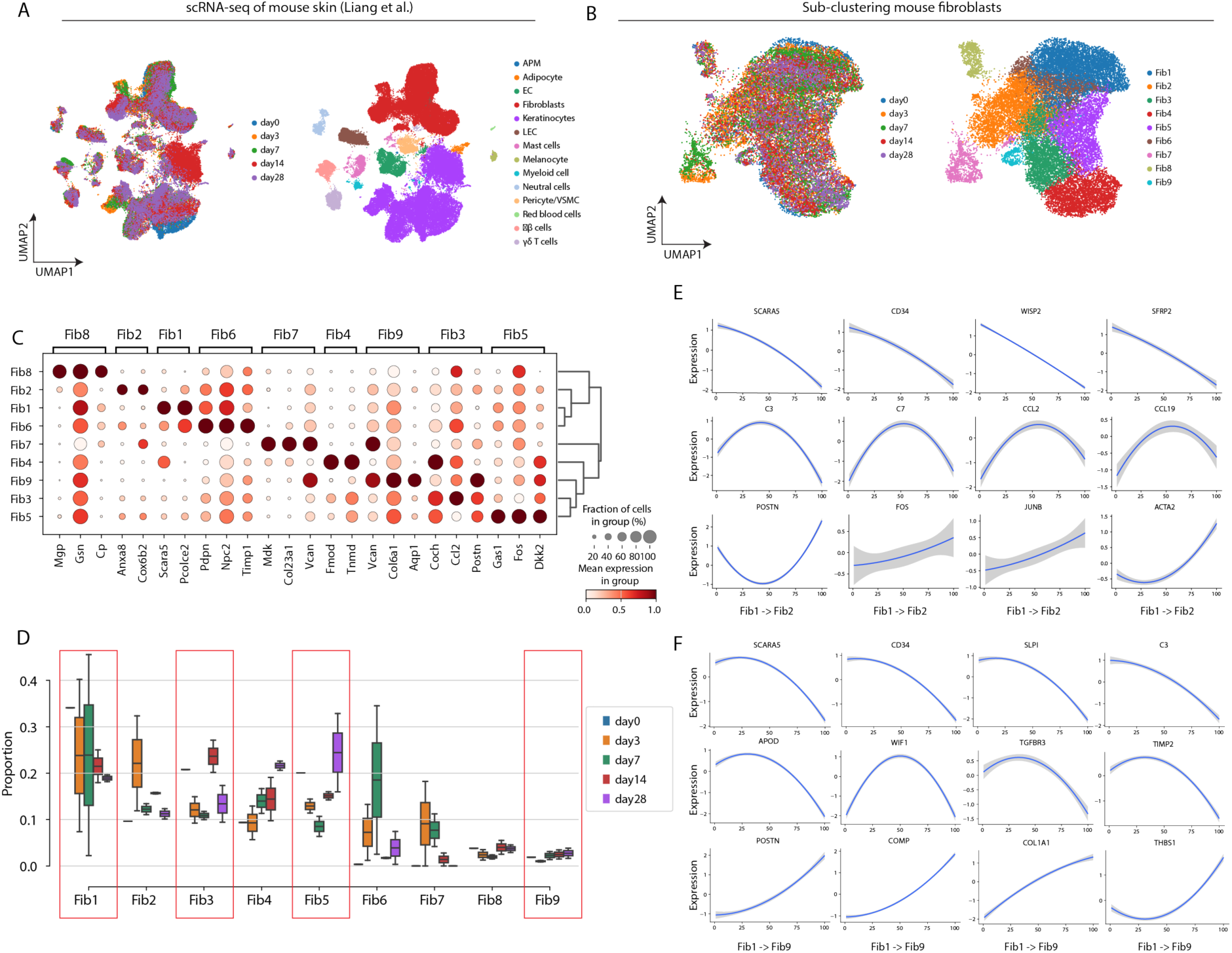
Trajectory analysis of fibroblast differentiation. (**A**), UMAP of scRNA-seq data from mouse skin(*26*) colored by time points (left) after bleomycin (BLM) treatment and annotated cell types (right). (**B**), UMAP of fibroblasts colored by time points (left) and sub-clusters (right). (**C**), Dot plots of marker genes for each sub-cluster. The colors refer to the mean expression of the gene, and the size represents the fraction of cells that expressed the gene. (**D**), Box plot showing the fraction of each sub-cluster within different time points. (**E**), Line plot showing expression dynamics of selected genes along the differentiation trajectory from Fib1 to Fib2. (F), Line plot showing expression dynamics of selected genes along the differentiation trajectory from Fib1 to Fib9.

**Fig. S7.**
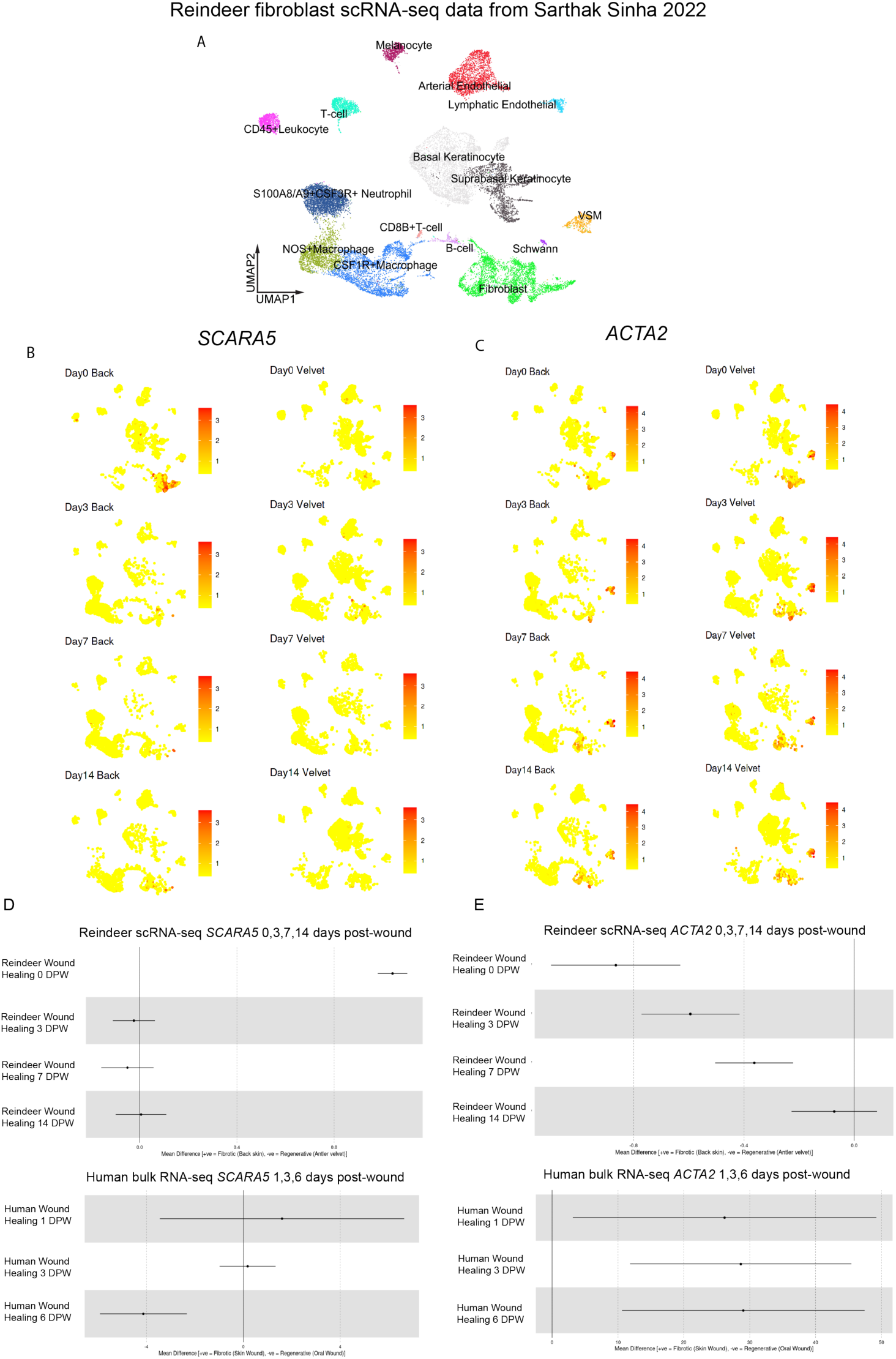
Cross-Species Validation of Fibroblast Dynamics. Sarthak Sinha et al. utilized single-cell multi-omics to investigate full-thickness skin injuries in reindeer, revealing that their antler skin (velvet) regenerates, while their back skin forms fibrotic scars(*7*). (**A**), UMAP of scRNA-seq data from reindeer skin colored by cell type. (**B**, **C**), Visualization of *SCARA5* (b) and *ACTA2* (c) expression in UMAP, showing the gene expression on the back (left) and on the velvet (right). (**D**, **E**), Line plot showing expression dynamics of *SCARA5* and *ACTA2* at the indicated days post wound in reindeer.

**Fig. S8.**
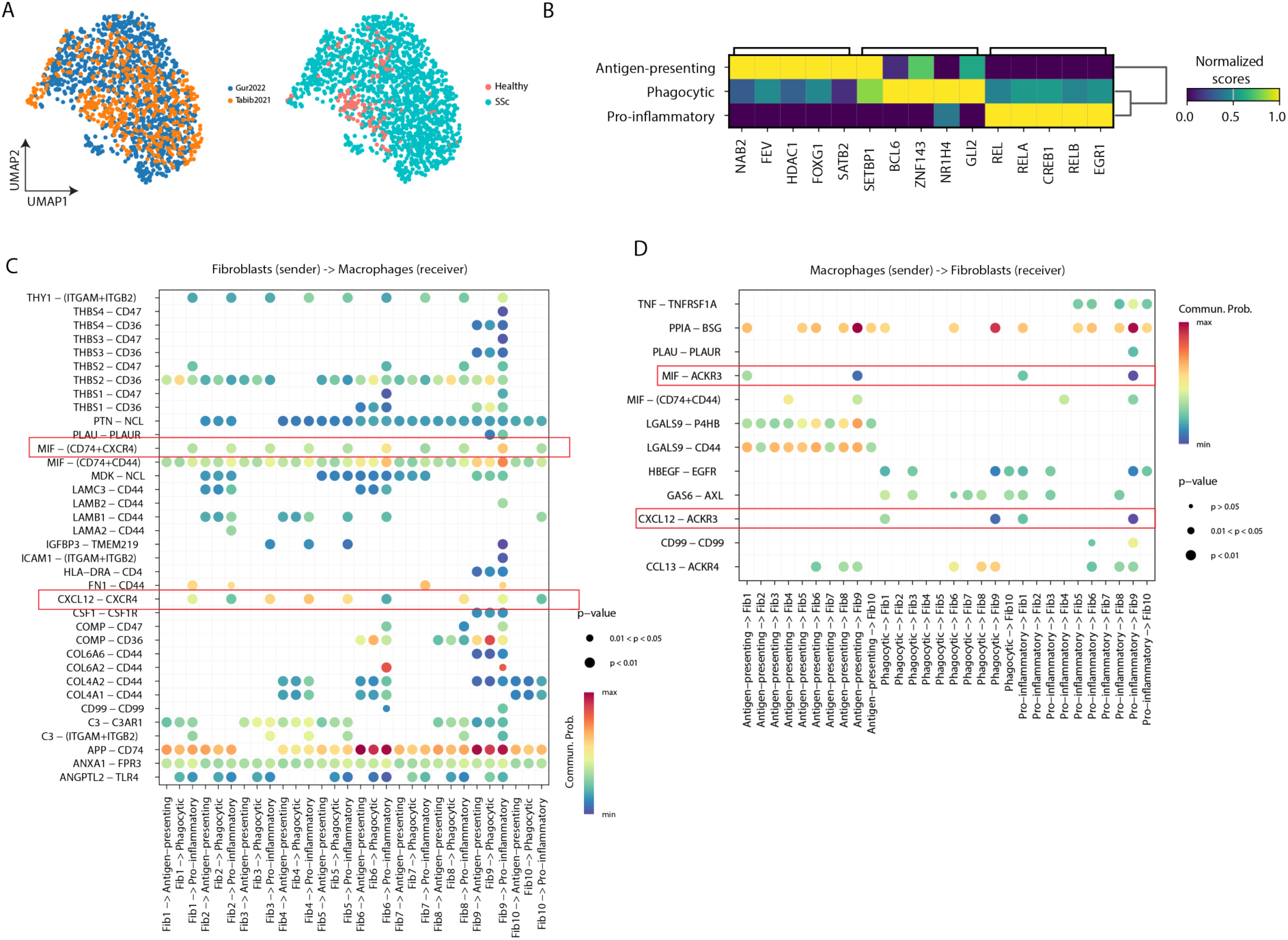
Sub-clustering of macrophages. (**A**), UMAP showing scRNA-seq of macrophages colored different datasets (left) and conditions (right). (**B**), Heamap of top four marker TFs for each sub-cluster-based TF activity score. (**C**), Heatmap showing all significant L-R pairs from fibroblasts to macrophages. (**D**), Heatmap showing all significant L-R pairs from macrophages to fibroblasts.

**Table. S1.** Clinical characterization of the patients with SSc undergoing skin biopsy for spatial transcriptomic study. * as categorized by the duration from first non-RP symptoms (early ≤ 36 months; late > 36 months); † as referred to the total mRSS assessed at 17 body sites; ‡ as defined by increase in total mRSS ≥ 25% from baseline to the follow-up visit within 52 weeks; || as detected by lung high-resolution CT; # as assessed by B-MODE Doppler echocardiography.

| Characteristics | N=10 |
| --- | --- |
| Gender (female/male) | 4/6 (40%/60%) |
| Median age at biopsy (range), years | 56.0 (27-73) |
| Subsets (dcSSc/lcSSc) | 10/0<br>(100%/0%) |
| Median disease duration from first non-RP (range), months | 8 (4-48) |
| Disease duration (early/late) * | 10/0<br>(100%/0%) |
| Median total mRSS (range) † | 21 (8-41) |
| Progression of skin fibrosis with increases in total mRSS ‡ | 10/0<br>(100%/0%) |
| Active digital ulcer (s) | 3 (30%) |
| Interstitial lung disease \|\| | 4 (40%) |
| Pulmonary arterial hypertension | 2 (20%) |
| Estimated systolic pulmonary artery pressure (sPAP) >30 mm Hg<sup>#</sup> | 3 (30%) |
| CRP (mg/dL) | 9.30<br>(1.49-45.38) |
| Disease specific autoantibodies |  |
| ATA positive | 3 (30%) |
| ARA positive | 4 (40%) |
| Anti-nRNP | 1 (10%) |
| Anti-Ku | 1 (10%) |
| Laboratory features |  |
| Hemoglobin (g/L) (Median, IQR) | 116.5,<br>100.0-136.0 |
| Leukocytes (10<sup>3</sup>/ul) (Median, IQR) | 6.46,<br>4.11-14.99 |
| Platelets (10<sup>3</sup>/ul) (Median, IQR) | 228, 120-363 |
| CRP (mg/l) (Median, IQR) | 4.60,<br>2.21-12.65 |

**Table. S2.** Quality statistics for 10x Visium and Stereo-seq data.

| sample | n_spot | condition | median_n_genes | median_n_UMIs | protocol |
| --- | --- | --- | --- | --- | --- |
| HC01 | 2035 | Healthy | 927 | 1170 | Visium |
| HC02 | 663 | Healthy | 1861 | 2517 | Visium |
| HC03 | 1805 | Healthy | 4096 | 8438 | Visium |
| HC05 | 827 | Healthy | 2352 | 3341 | Visium |
| SSc-HL01 | 453 | SSc | 3477 | 6437 | Visium |
| SSc-HL05 | 508 | SSc | 2864 | 5895.5 | Visium |
| SSc-HL06 | 612 | SSc | 3253.5 | 5587.5 | Visium |
| SSc-HL11 | 695 | SSc | 1266 | 1850 | Visium |
| SSc-HL13 | 605 | SSc | 1056 | 1495 | Visium |
| SSc-HL25 | 751 | SSc | 1757 | 2935 | Visium |
| SSc-HL33 | 737 | SSc | 2247 | 3629 | Visium |
| SSc-HL35 | 877 | SSc | 3087 | 5945 | Visium |
| SSc4994 | 1745 | SSc | 2238 | 3575 | Visium |
| SSc5380 | 891 | SSc | 1482 | 1980 | Visium |
| HC01 | 17089 | Healthy | 212 | 272 | Stereo-seq |
| HL05 | 3437 | SSc | 330 | 440 | Stereo-seq |
| HL25 | 10928 | SSc | 226 | 289 | Stereo-seq |
| HL35 | 9112 | SSc | 184 | 234 | Stereo-seq |

## References

1. M. Elhai, C. Meune, M. Boubaya, J. Avouac, E. Hachulla, A. Balbir-Gurman, G. Riemekasten, P. Airò, B. Joven, S. Vettori, F. Cozzi, S. Ullman, L. Czirják, M. Tikly, U. Müller-Ladner, P. Caramaschi, O. Distler, F. Iannone, L. P. Ananieva, R. Hesselstrand, R. Becvar, A. Gabrielli, N. Damjanov, M. J. Salvador, V. Riccieri, C. Mihai, G. Szücs, U. A. Walker, N. Hunzelmann, D. Martinovic, V. Smith, C. de S. Müller, C. M. Montecucco, D. Opris, F. Ingegnoli, P. G. Vlachoyiannopoulos, B. Stamenkovic, E. Rosato, S. Heitmann, J. H. W. Distler, T. Zenone, M. Seidel, A. Vacca, E. D. Langhe, S. Novak, M. Cutolo, L. Mouthon, J. Henes, C. Chizzolini, C. A. von Mühlen, K. Solanki, S. Rednic, L. Stamp, B. Anic, V. O. Santamaria, M. De Santis, S. Yavuz, W. A. Sifuentes-Giraldo, E. Chatelus, J. Stork, J. van Laar, E. Loyo, P. García de la Peña Lefebvre, K. Eyerich, V. Cosentino, J. J. Alegre-Sancho, O. Kowal-Bielecka, G. Rey, M. Matucci-Cerinic, Y. Allanore, EUSTAR group, Mapping and predicting mortality from systemic sclerosis. Ann. Rheum. Dis. 76, 1897–1905 (2017).

2. J. H. W. Distler, A.-H. Györfi, M. Ramanujam, M. L. Whitfield, M. Königshoff, R. Lafyatis, Shared and distinct mechanisms of fibrosis. Nat. Rev. Rheumatol. 15, 705–730 (2019).

3. M. Alexanian, A. Padmanabhan, T. Nishino, J. G. Travers, L. Ye, A. Pelonero, C. Y. Lee, N. Sadagopan, Y. Huang, K. Auclair, A. Zhu, Y. An, C. A. Ekstrand, C. Martinez, B. G. Teran, W. R. Flanigan, C. K.-S. Kim, K. Lumbao-Conradson, Z. Gardner, L. Li, M. W. Costa, R. Jain, I. Charo, A. J. Combes, S. M. Haldar, K. S. Pollard, R. J. Vagnozzi, T. A. McKinsey, P. F. Przytycki, D. Srivastava, Chromatin remodelling drives immune cell-fibroblast communication in heart failure. Nature 635, 434–443 (2024).

4. J. M. Amrute, X. Luo, V. Penna, S. Yang, T. Yamawaki, S. Hayat, A. Bredemeyer, I.-H. Jung, F. F. Kadyrov, G. S. Heo, R. Venkatesan, S. Y. Shi, A. Parvathaneni, A. L. Koenig, C. Kuppe, C. Baker, H. Luehmann, C. Jones, B. Kopecky, X. Zeng, T. Bleckwehl, P. Ma, P. Lee, Y. Terada, A. Fu, M. Furtado, D. Kreisel, A. Kovacs, N. O. Stitziel, S. Jackson, C.-M. Li, Y. Liu, N. A. Rosenthal, R. Kramann, B. Ason, K. J. Lavine, Targeting immune-fibroblast cell communication in heart failure. Nature 635, 423–433 (2024).

5. C. H. Mayr, D. Santacruz, S. Jarosch, M. Bleck, J. Dalton, A. McNabola, C. Lempp, L. Neubert, B. Rath, J. C. Kamp, D. Jonigk, M. Kühnel, H. Schlüter, A. Klimowicz, J. Doerr, A. Dick, F. Ramirez, M. J. Thomas, Spatial transcriptomic characterization of pathologic niches in IPF. Sci. Adv. 10, eadl5473 (2024).

6. C. Morse, T. Tabib, J. Sembrat, K. L. Buschur, H. T. Bittar, E. Valenzi, Y. Jiang, D. J. Kass, K. Gibson, W. Chen, A. Mora, P. V. Benos, M. Rojas, R. Lafyatis, Proliferating SPP1/MERTK-expressing macrophages in idiopathic pulmonary fibrosis. Eur. Respir. J. 54, 1802441 (2019).

7. S. Sinha, H. D. Sparks, E. Labit, H. N. Robbins, K. Gowing, A. Jaffer, E. Kutluberk, R. Arora, M. S. B. Raredon, L. Cao, S. Swanson, P. Jiang, O. Hee, H. Pope, M. Workentine, K. Todkar, N. Sharma, S. Bharadia, K. Chockalingam, L. G. N. de Almeida, M. Adam, L. Niklason, S. S. Potter, A. W. Seifert, A. Dufour, V. Gabriel, N. L. Rosin, R. Stewart, G. Muench, R. McCorkell, J. Matyas, J. Biernaskie, Fibroblast inflammatory priming determines regenerative versus fibrotic skin repair in reindeer. Cell 185, 4717–4736.e25 (2022).

8. S. Chakarov, H. Y. Lim, L. Tan, S. Y. Lim, P. See, J. Lum, X.-M. Zhang, S. Foo, S. Nakamizo, K. Duan, W. T. Kong, R. Gentek, A. Balachander, D. Carbajo, C. Bleriot, B. Malleret, J. K. C. Tam, S. Baig, M. Shabeer, S.-A. E. S. Toh, A. Schlitzer, A. Larbi, T. Marichal, B. Malissen, J. Chen, M. Poidinger, K. Kabashima, M. Bajenoff, L. G. Ng, V. Angeli, F. Ginhoux, Two distinct interstitial macrophage populations coexist across tissues in specific subtissular niches. Science 363, eaau0964 (2019).

9. T. Tabib, M. Huang, N. Morse, A. Papazoglou, R. Behera, M. Jia, M. Bulik, D. E. Monier, P. V. Benos, W. Chen, R. Domsic, R. Lafyatis, Myofibroblast transcriptome indicates SFRP2hi fibroblast progenitors in systemic sclerosis skin. Nat. Commun. 12, 4384 (2021).

10. C. Gur, S.-Y. Wang, F. Sheban, M. Zada, B. Li, F. Kharouf, H. Peleg, S. Aamar, A. Yalin, D. Kirschenbaum, Y. Braun-Moscovici, D. A. Jaitin, T. Meir-Salame, E. Hagai, B. K. Kragesteen, B. Avni, S. Grisariu, C. Bornstein, S. Shlomi-Loubaton, E. David, R. Shreberk-Hassidim, V. Molho-Pessach, D. Amar, T. Tzur, R. Kuint, M. Gross, O. Barboy, A. Moshe, L. Fellus-Alyagor, D. Hirsch, Y. Addadi, S. Erenfeld, M. Biton, T. Tzemach, A. Elazary, Y. Naparstek, R. Tzemach, A. Weiner, A. Giladi, A. Balbir-Gurman, I. Amit, LGR5 expressing skin fibroblasts define a major cellular hub perturbed in scleroderma. Cell 185, 1373–1388.e20 (2022).

11. K. E. N. Clark, S. Xu, M. Attah, V. H. Ong, C. D. Buckley, C. P. Denton, Single-cell analysis reveals key differences between early-stage and late-stage systemic sclerosis skin across autoantibody subgroups. Ann. Rheum. Dis. 82, 1568–1579 (2023).

12. H. Yin, O. Distler, L. Shen, X. Xu, Y. Yuan, R. Li, B. Liu, Q. Li, Q. Huang, F. Xie, Z. Zhang, R. Liang, X. Dai, X. Chen, B. Li, Q. Yan, L. Lu, Endothelial response to type I interferon contributes to vasculopathy and fibrosis and predicts disease progression of systemic sclerosis. Arthritis Rheumatol. 76, 78–91 (2024).

13. F. Ma, P.-S. Tsou, M. Gharaee-Kermani, O. Plazyo, X. Xing, J. Kirma, R. Wasikowski, G. A. Hile, P. W. Harms, Y. Jiang, E. Xing, M. Nakamura, D. Ochocki, W. D. Brodie, S. Pillai, E. Maverakis, M. Pellegrini, R. L. Modlin, J. Varga, L. C. Tsoi, R. Lafyatis, J. M. Kahlenberg, A. C. Billi, D. Khanna, J. E. Gudjonsson, Systems-based identification of the Hippo pathway for promoting fibrotic mesenchymal differentiation in systemic sclerosis. Nat. Commun. 15, 210 (2024).

14. A. Rius Rigau, M. Liang, V. Devakumar, R. Neelagar, A.-E. Matei, A.-H. Györfi, C. Bergmann, T. Filla, V. Fedorchenko, G. Schett, J. H. W. Distler, Y.-N. Li, Imaging mass cytometry-based characterisation of fibroblast subsets and their cellular niches in systemic sclerosis. Ann. Rheum. Dis., doi: 10.1136/ard-2024-226336 (2024).

15. A. Rius Rigau, Y.-N. Li, A.-E. Matei, A.-H. Györfi, P.-M. Bruch, S. Koziel, V. Devakumar, A. Gabrielli, A. Kreuter, J. Wang, S. Dietrich, G. Schett, J. H. W. Distler, M. Liang, Characterization of vascular niche in systemic sclerosis by spatial proteomics. Circ. Res. 134, 875–891 (2024).

16. A. Chen, S. Liao, M. Cheng, K. Ma, L. Wu, Y. Lai, X. Qiu, J. Yang, J. Xu, S. Hao, X. Wang, H. Lu, X. Chen, X. Liu, X. Huang, Z. Li, Y. Hong, Y. Jiang, J. Peng, S. Liu, M. Shen, C. Liu, Q. Li, Y. Yuan, X. Wei, H. Zheng, W. Feng, Z. Wang, Y. Liu, Z. Wang, Y. Yang, H. Xiang, L. Han, B. Qin, P. Guo, G. Lai, P. Muñoz-Cánoves, P. H. Maxwell, J. P. Thiery, Q.-F. Wu, F. Zhao, B. Chen, M. Li, X. Dai, S. Wang, H. Kuang, J. Hui, L. Wang, J.-F. Fei, O. Wang, X. Wei, H. Lu, B. Wang, S. Liu, Y. Gu, M. Ni, W. Zhang, F. Mu, Y. Yin, H. Yang, M. Lisby, R. J. Cornall, J. Mulder, M. Uhlén, M. A. Esteban, Y. Li, L. Liu, X. Xu, J. Wang, Spatiotemporal transcriptomic atlas of mouse organogenesis using DNA nanoball-patterned arrays. Cell 185, 1777–1792.e21 (2022).

17. J. Avouac, K. Palumbo, M. Tomcik, P. Zerr, C. Dees, A. Horn, B. Maurer, A. Akhmetshina, C. Beyer, A. Sadowski, H. Schneider, S. Shiozawa, O. Distler, G. Schett, Y. Allanore, J. H. W. Distler, Inhibition of activator protein 1 signaling abrogates transforming growth factor β-mediated activation of fibroblasts and prevents experimental fibrosis. Arthritis Rheum. 64, 1642–1652 (2012).

18. C. Bergmann, S. Chenguiti Fakhouri, T.-M. Thuong, T. Filla, A. R. Rigau, A. B. Ekici, B. Merlevede, L. Hallenberger, H. Zhu, C. Dees, A.-E. Matei, J. Auth, A.-H. Györfi, X. Zhou, S. Rauber, A. Bozec, N. Dickel, C. Liang, M. Kunz, G. Schett, J. H. W. Distler, Mutual amplification of GLI2/Hedgehog and cJUN/AP1 signaling in fibroblasts in Systemic Sclerosis (SSc) - potential implications for combined therapies. Arthritis Rheumatol., doi: 10.1002/art.42979 (2024).

19. C. Kuppe, M. M. Ibrahim, J. Kranz, X. Zhang, S. Ziegler, J. Perales-Patón, J. Jansen, K. C. Reimer, J. R. Smith, R. Dobie, J. R. Wilson-Kanamori, M. Halder, Y. Xu, N. Kabgani, N. Kaesler, M. Klaus, L. Gernhold, V. G. Puelles, T. B. Huber, P. Boor, S. Menzel, R. M. Hoogenboezem, E. M. J. Bindels, J. Steffens, J. Floege, R. K. Schneider, J. Saez-Rodriguez, N. C. Henderson, R. Kramann, Decoding myofibroblast origins in human kidney fibrosis. Nature 589, 281–286 (2021).

20. C. Kuppe, R. O. Ramirez Flores, Z. Li, S. Hayat, R. T. Levinson, X. Liao, M. T. Hannani, J. Tanevski, F. Wünnemann, J. S. Nagai, M. Halder, D. Schumacher, S. Menzel, G. Schäfer, K. Hoeft, M. Cheng, S. Ziegler, X. Zhang, F. Peisker, N. Kaesler, T. Saritas, Y. Xu, A. Kassner, J. Gummert, M. Morshuis, J. Amrute, R. J. A. Veltrop, P. Boor, K. Klingel, L. W. Van Laake, A. Vink, R. M. Hoogenboezem, E. M. J. Bindels, L. Schurgers, S. Sattler, D. Schapiro, R. K. Schneider, K. Lavine, H. Milting, I. G. Costa, J. Saez-Rodriguez, R. Kramann, Spatial multi-omic map of human myocardial infarction. Nature 608, 766–777 (2022).

21. Z. Li, C. Kuppe, S. Ziegler, M. Cheng, N. Kabgani, S. Menzel, M. Zenke, R. Kramann, I. G. Costa, Chromatin-accessibility estimation from single-cell ATAC-seq data with scOpen. Nat. Commun. 12, 6386 (2021).

22. Y. Nanri, S. Nunomura, Y. Terasaki, T. Yoshihara, Y. Hirano, Y. Yokosaki, Y. Yamaguchi, C. Feghali-Bostwick, K. Ajito, S. Murakami, S. J. Conway, K. Izuhara, Cross-talk between transforming growth factor-β and periostin can be targeted for pulmonary fibrosis. Am. J. Respir. Cell Mol. Biol. 62, 204–216 (2020).

23. Y. Nanri, S. Nunomura, Y. Honda, H. Takedomi, Y. Yamaguchi, K. Izuhara, A positive loop formed by SOX11 and periostin upregulates TGF-β signals leading to skin fibrosis. J. Invest. Dermatol. 143, 989–998.e7 (2023).

24. R. Liang, B. Šumová, C. Cordazzo, T. Mallano, Y. Zhang, T. Wohlfahrt, C. Dees, A. Ramming, D. Krasowska, M. Michalska-Jakubus, O. Distler, G. Schett, L. Šenolt, J. H. W. Distler, The transcription factor GLI2 as a downstream mediator of transforming growth factor-β-induced fibroblast activation in SSc. Ann. Rheum. Dis. 76, 756–764 (2017).

25. B. Skaug, D. Khanna, W. R. Swindell, M. E. Hinchcliff, T. M. Frech, V. D. Steen, F. N. Hant, J. K. Gordon, A. A. Shah, L. Zhu, W. J. Zheng, J. L. Browning, A. M. S. Barron, M. Wu, S. Visvanathan, P. Baum, J. M. Franks, M. L. Whitfield, V. K. Shanmugam, R. T. Domsic, F. V. Castelino, E. J. Bernstein, N. Wareing, M. A. Lyons, J. Ying, J. Charles, M. D. Mayes, S. Assassi, Global skin gene expression analysis of early diffuse cutaneous systemic sclerosis shows a prominent innate and adaptive inflammatory profile. Ann. Rheum. Dis. 79, 379–386 (2020).

26. Y. Liang, Y. Hu, J. Zhang, H. Song, X. Zhang, Y. Chen, Y. Peng, L. Sun, Y. Sun, R. Xue, S. Ji, C. Li, Z. Rong, B. Yang, Y. Xu, Dynamic pathological analysis reveals a protective role against skin fibrosis for TREM2-dependent macrophages. Theranostics 14, 2232– 2245 (2024).

27. B. Nazari, L. M. Rice, G. Stifano, A. M. S. Barron, Y. M. Wang, T. Korndorf, J. Lee, J. Bhawan, R. Lafyatis, J. L. Browning, Altered dermal fibroblasts in systemic sclerosis display podoplanin and CD90. Am. J. Pathol. 186, 2650–2664 (2016).

28. J. Tanevski, R. O. R. Flores, A. Gabor, D. Schapiro, J. Saez-Rodriguez, Explainable multiview framework for dissecting spatial relationships from highly multiplexed data. Genome Biol. 23, 97 (2022).

29. D. Khanna, S. Huang, C. J. F, Lin, C. Spino. New Composite Endpoint Early Diffuse Cutaneous Systemic Sclerosis-Revisiting Provisional American College Rheumatology Composite Response Index Systemic Sclerosis.Ann Rheum Dis. 80, 641–650 (2021).

30. M. B. Buechler, W. Fu, S. J. Turley, Fibroblast-macrophage reciprocal interactions in health, fibrosis, and cancer. Immunity 54, 903–915 (2021).

31. R. Bhandari, M. S. Ball, V. Martyanov, D. Popovich, E. Schaafsma, S. Han, M. ElTanbouly, N. M. Orzechowski, M. Carns, E. Arroyo, K. Aren, M. Hinchcliff, M. L. Whitfield, P. A. Pioli, Profibrotic activation of human macrophages in systemic sclerosis. Arthritis Rheumatol. 72, 1160–1169 (2020).

32. S. Boulet, L. Le Corre, L. Odagiu, N. Labrecque, Role of NR4A family members in myeloid cells and leukemia. Curr. Res. Immunol. 3, 23–36 (2022).

33. K. Palumbo-Zerr, P. Zerr, A. Distler, J. Fliehr, R. Mancuso, J. Huang, D. Mielenz, M. Tomcik, B. G. Fürnrohr, C. Scholtysek, C. Dees, C. Beyer, G. Krönke, D. Metzger, O. Distler, G. Schett, J. H. W. Distler, Orphan nuclear receptor NR4A1 regulates transforming growth factor-β signaling and fibrosis. Nat. Med. 21, 150–158 (2015).

34. D. Kwak, P. B. Bradley, N. Subbotina, S. Ling, S. Teitz-Tennenbaum, J. J. Osterholzer, T. H. Sisson, K. K. Kim, CD36/Lyn kinase interactions within macrophages promotes pulmonary fibrosis in response to oxidized phospholipid. Respir. Res. 24, 314 (2023).

35. S. Pennathur, K. Pasichnyk, N. M. Bahrami, L. Zeng, M. Febbraio, I. Yamaguchi, D. M. Okamura, The macrophage phagocytic receptor CD36 promotes fibrogenic pathways on removal of apoptotic cells during chronic kidney injury. Am. J. Pathol. 185, 2232–2245 (2015).

36. A. Papazoglou, M. Huang, M. Bulik, A. Lafyatis, T. Tabib, C. Morse, J. Sembrat, M. Rojas, E. Valenzi, R. Lafyatis, Epigenetic regulation of profibrotic macrophages in systemic sclerosis-associated interstitial lung disease. Arthritis Rheumatol. 74, 2003–2014 (2022).

37. D. Xu, S. Bhattacharyya, W. Wang, I. Ifergan, M.-Y. A. C. Wong, D. Procissi, A. Yeldandi, S. Bale, R. G. Marangoni, C. Horbinski, S. D. Miller, J. Varga, PLG nanoparticles target fibroblasts and MARCO+ monocytes to reverse multiorgan fibrosis. JCI Insight 7 (2022).

38. J. E. Anstee, K. T. Feehan, J. W. Opzoomer, I. Dean, H. P. Muller, M. Bahri, T. S. Cheung, K. Liakath-Ali, Z. Liu, D. Choy, J. Caron, D. Sosnowska, R. Beatson, T. Muliaditan, Z. An, C. E. Gillett, G. Lan, X. Zou, F. M. Watt, T. Ng, J. M. Burchell, S. Kordasti, D. R. Withers, T. Lawrence, J. N. Arnold, LYVE-1+ macrophages form a collaborative CCR5-dependent perivascular niche that influences chemotherapy responses in murine breast cancer. Dev. Cell 58, 1548–1561.e10 (2023).

39. T. Liu, L. Zhang, D. Joo, S.-C. Sun, NF-κB signaling in inflammation. Signal Transduct. Target. Ther. 2 (2017).

40. S. Jin, M. V. Plikus, Q. Nie, CellChat for systematic analysis of cell-cell communication from single-cell transcriptomics. Nat. Protoc., doi: 10.1038/s41596-024-01045-4 (2024).

41. G. Cuesta-Margolles, G. Schlecht-Louf, F. Bachelerie, ACKR3 in skin homeostasis, an overlooked player in the CXCR4/CXCL12 axis. J. Invest. Dermatol., doi: 10.1016/j.jid.2024.08.022 (2024).

42. F. Schreibing, T. M. Anslinger, R. Kramann, Fibrosis in pathology of heart and kidney: From deep RNA-sequencing to novel molecular targets. Circ. Res. 132, 1013–1033 (2023).

43. Y. Gao, J. Li, W. Cheng, T. Diao, H. Liu, Y. Bo, C. Liu, W. Zhou, M. Chen, Y. Zhang, Z. Liu, W. Han, R. Chen, J. Peng, L. Zhu, W. Hou, Z. Zhang, Cross-tissue human fibroblast atlas reveals myofibroblast subtypes with distinct roles in immune modulation. Cancer Cell 42, 1764–1783.e10 (2024).

44. K. R. Flaherty, A. U. Wells, V. Cottin, A. Devaraj, S. L. F. Walsh, Y. Inoue, L. Richeldi, M. Kolb, K. Tetzlaff, S. Stowasser, C. Coeck, E. Clerisme-Beaty, B. Rosenstock, M. Quaresma, T. Haeufel, R.-G. Goeldner, R. Schlenker-Herceg, K. K. Brown, INBUILD Trial Investigators, Nintedanib in progressive fibrosing interstitial lung diseases. N. Engl. J. Med. 381, 1718–1727 (2019).

45. C. Grönberg, S. Rattik, C. Tran-Manh, X. Zhou, A. Rius Rigau, Y.-N. Li, A.-H. Györfi, N. Dickel, M. Kunz, A. Kreuter, E.-A. Matei, H. Zhu, P. Skoog, D. Liberg, J. H. Distler, T. Trinh-Minh, Combined inhibition of IL-1, IL-33 and IL-36 signalling by targeting IL1RAP ameliorates skin and lung fibrosis in preclinical models of systemic sclerosis. Ann. Rheum. Dis. 83, 1156–1168 (2024).

46. F. van den Hoogen, D. Khanna, J. Fransen, S. R. Johnson, M. Baron, A. Tyndall, M. Matucci-Cerinic, R. P. Naden, T. A. Medsger Jr, P. E. Carreira, G. Riemekasten, P. J. Clements, C. P. Denton, O. Distler, Y. Allanore, D. E. Furst, A. Gabrielli, M. D. Mayes, J. M. van Laar, J. R. Seibold, L. Czirjak, V. D. Steen, M. Inanc, O. Kowal-Bielecka, U. Müller-Ladner, G. Valentini, D. J. Veale, M. C. Vonk, U. A. Walker, L. Chung, D. H. Collier, M. E. Csuka, B. J. Fessler, S. Guiducci, A. Herrick, V. M. Hsu, S. Jimenez, B. Kahaleh, P. A. Merkel, S. Sierakowski, R. M. Silver, R. W. Simms, J. Varga, J. E. Pope, 2013 classification criteria for systemic sclerosis: an American College of Rheumatology/European League against Rheumatism collaborative initiative: ACR/EULAR classification criteria for SSc. Arthritis Rheum. 65, 2737–2747 (2013).

47. A. Chen, Y. Sun, Y. Lei, C. Li, S. Liao, J. Meng, Y. Bai, Z. Liu, Z. Liang, Z. Zhu, N. Yuan, H. Yang, Z. Wu, F. Lin, K. Wang, M. Li, S. Zhang, M. Yang, T. Fei, Z. Zhuang, Y. Huang, Y. Zhang, Y. Xu, L. Cui, R. Zhang, L. Han, X. Sun, B. Chen, W. Li, B. Huangfu, K. Ma, J. Ma, Z. Li, Y. Lin, H. Wang, Y. Zhong, H. Zhang, Q. Yu, Y. Wang, X. Liu, J. Peng, C. Liu, W. Chen, W. Pan, Y. An, S. Xia, Y. Lu, M. Wang, X. Song, S. Liu, Z. Wang, C. Gong, X. Huang, Y. Yuan, Y. Zhao, Q. Chai, X. Tan, J. Liu, M. Zheng, S. Li, Y. Huang, Y. Hong, Z. Huang, M. Li, M. Jin, Y. Li, H. Zhang, S. Sun, L. Gao, Y. Bai, M. Cheng, G. Hu, S. Liu, B. Wang, B. Xiang, S. Li, H. Li, M. Chen, S. Wang, M. Li, W. Liu, X. Liu, Q. Zhao, M. Lisby, J. Wang, J. Fang, Y. Lin, Q. Xie, Z. Liu, J. He, H. Xu, W. Huang, J. Mulder, H. Yang, Y. Sun, M. Uhlen, M. Poo, J. Wang, J. Yao, W. Wei, Y. Li, Z. Shen, L. Liu, Z. Liu, X. Xu, C. Li, Single-cell spatial transcriptome reveals cell-type organization in the macaque cortex. Cell 186, 3726–3743.e24 (2023).

48. F. A. Wolf, P. Angerer, F. J. Theis, SCANPY: large-scale single-cell gene expression data analysis. Genome Biol. 19, 15 (2018).

49. I. Korsunsky, N. Millard, J. Fan, K. Slowikowski, F. Zhang, K. Wei, Y. Baglaenko, M. Brenner, P.-R. Loh, S. Raychaudhuri, Fast, sensitive and accurate integration of single-cell data with Harmony. Nat. Methods 16, 1289–1296 (2019).

50. V. Kleshchevnikov, A. Shmatko, E. Dann, A. Aivazidis, H. W. King, T. Li, R. Elmentaite, A. Lomakin, V. Kedlian, A. Gayoso, M. S. Jain, J. S. Park, L. Ramona, E. Tuck, A. Arutyunyan, R. Vento-Tormo, M. Gerstung, L. James, O. Stegle, O. A. Bayraktar, Cell2location maps fine-grained cell types in spatial transcriptomics. Nat. Biotechnol. 40, 661–671 (2022).

51. P. Badia-I-Mompel, J. Vélez Santiago, J. Braunger, C. Geiss, D. Dimitrov, S. Müller-Dott, P. Taus, A. Dugourd, C. H. Holland, R. O. Ramirez Flores, J. Saez-Rodriguez, decoupleR: ensemble of computational methods to infer biological activities from omics data. Bioinform Adv 2, vbac016 (2022).

52. V. A. Traag, L. Waltman, N. J. van Eck, From Louvain to Leiden: guaranteeing well-connected communities. Sci. Rep. 9, 5233 (2019).

53. Y. Hao, T. Stuart, M. H. Kowalski, S. Choudhary, P. Hoffman, A. Hartman, A. Srivastava, G. Molla, S. Madad, C. Fernandez-Granda, R. Satija, Dictionary learning for integrative, multimodal and scalable single-cell analysis. Nat. Biotechnol., doi: 10.1038/s41587-023-01767-y (2023).

54. J. M. Granja, M. R. Corces, S. E. Pierce, S. T. Bagdatli, H. Choudhry, H. Y. Chang, W. J. Greenleaf, ArchR is a scalable software package for integrative single-cell chromatin accessibility analysis. Nat. Genet. 53, 403–411 (2021).

55. M. Liang, L. Wang, X. Tian, K. Wang, X. Zhu, L. Huang, Q. Li, W. Ye, C. Chen, H. Yang, W. Wu, X. Chen, X. Zhu, Y. Xue, W. Wan, Y. Wu, L. Lu, J. Wang, H. Zou, T. Ying, F. Zhou, Identification and validation of anti-protein arginine methyltransferase 5 (PRMT5) antibody as a novel biomarker for systemic sclerosis (SSc). Ann. Rheum. Dis. 83, 1144– 1155 (2024).

